# Mechanistic basis for ubiquitin modulation of a protein energy landscape

**DOI:** 10.1101/2020.12.06.414011

**Authors:** Emma C. Carroll, Naomi R. Latorraca, Johanna M. Lindner, Brendan C. Maguire, Jeff G. Pelton, Susan Marqusee

## Abstract

Ubiquitin is a common posttranslational modification canonically associated with targeting proteins to the 26S proteasome for degradation and also plays a role in numerous other non-degradative cellular processes. Ubiquitination at certain sites destabilizes the substrate protein, with consequences for proteasomal processing, while ubiquitination at other sites has little energetic effect. How this site specificity—and, by extension, the myriad effects of ubiquitination on substrate proteins—arises remains unknown. Here, we systematically characterize the atomic-level effects of ubiquitination at various sites on a model protein, barstar, using a combination of NMR, hydrogen-deuterium exchange mass spectrometry, and molecular dynamics simulation. We find that, regardless of the site of modification, ubiquitination does not induce large structural rearrangements in the substrate. Destabilizing modifications, however, increase fluctuations from the native state resulting in exposure of the substrate’s C terminus. Both of the sites occur in regions of barstar with relatively high conformational flexibility. Destabilization, however, appears to occur through different thermodynamic mechanisms, involving a reduction in entropy in one case and a loss in enthalpy in another. By contrast, ubiquitination at a non-destabilizing site protects the substrate C terminus through intermittent formation of a structural motif with the last three residues of ubiquitin. Thus, the biophysical effects of ubiquitination at a given site depend greatly on local context. Taken together, our results reveal how a single post-translational modification can generate a broad array of distinct effects, providing a framework to guide the design of proteins and therapeutics with desired degradation and quality-control properties. (248 words)

**Significance Statement:** Fluctuations on a protein energy landscapes encode the mechanistic basis for vital biological processes not always evident from static structures alone. Ubiquitination, a key posttranslational modification, can affect a protein’s energy landscape with consequences for proteasomal degradation, but the molecular mechanisms driving ubiquitin-induced energetic changes remain elusive. Here, we systematically characterize the energetic effects of ubiquitination at three sites on a model protein. We find that distinct thermodynamic mechanisms can produce the same outcome of ubiquitin-induced destabilization at sensitive sites. At a non-sensitive site, we observe formation of a substrate–ubiquitin interaction that may serve to protect against destabilization. This work will enable development of predictive models of the effect of ubiquitin at any given site on a protein with implications for understanding and engineering regulated ubiquitin signaling and protein quality control in vivo.

## Introduction

Ubiquitin is an 8.5 kDa protein appended to target proteins as a posttranslational modification (PTM). Typically, ubiquitin is conjugated to the primary amine of substrate lysine residues, though non-canonical linkages to serine and cysteine also exist in vivo. Ubiquitin itself contains seven lysine residues, which allows building of ubiquitin chains with various linkages and topologies. Ubiquitination is most typically associated with targeting condemned proteins to the 26S proteasome for degradation; however, it is also involved in a large and ever-growing list of crucial regulatory, non-degradative cellular processes (1). A complex and highly regulated enzymatic cascade attaches ubiquitin to substrates and therefore plays a key role in determining the specific downstream effects of an individual ubiquitination event. There are several hundred E3 ligases, the terminal enzymes in this cascade (2), which give rise to broad proteome coverage and allow for some level of site specificity (3, 4).

Multiple different ubiquitin chain linkages and topologies bind with high affinity to proteasomal ubiquitin receptors and promote degradation (5–8). However, the presence of a ubiquitin tag alone is not sufficient to ensure proteasomal degradation. In fact, a substantial proportion of ubiquitin-modified proteins that interact with the 26S proteasome are ultimately released (9, 10) and not degraded. The proteasome also relies on substrate conformational properties, initiating degradation at an unstructured region on the condemned protein (11, 12). Much work has been done to understand the requirements of this unstructured region with regard to length, sequence composition, and topological position (13–15), yet at least 30% of known proteasome clients lack such a region (16). While evidence suggests that well-folded proteins are processed by diverse cellular unfoldases, such as Cdc48/p97/VCP (17, 18), an intriguing possibility is that the ubiquitin modification itself can modulate the conformational landscape and thus regulate proteasome substrate selection. Simulations have suggested that ubiquitination can destabilize the folded state of the substrate protein, thereby allowing it to more readily adopt unfolded or partially unfolded conformations (19, 20).

Recently, we demonstrated that this is indeed the case: ubiquitin can exert significant effects on a substrate’s energy landscape depending on the site of ubiquitination and the identity of the substrate protein. Moreover, these changes can have direct consequences for proteasomal processing (21). By examining the energetic effects of native, isopeptide-linked ubiquitin attachment to three different sites within the small protein barstar from *bacillus amyloliquefaciens*, we found that ubiquitin attached at either lysine 2 or lysine 60 destabilizes the protein both globally and via subglobal fluctuations, and we thus refer to these residues as sensitive sites. By contrast, ubiquitination at lysine 78 produces little effect on the energy landscape (21), and we therefore term it a non-sensitive site. Another study found that ubiquitin, appended through a non-native linkage, can destabilize a folded substrate as measured by changes in the midpoints for thermally induced unfolding transitions (22).

Ubiquitination at the two sensitive sites in barstar increases the population of partially unfolded, high-energy states on the landscape sufficient for proteasomal engagement and degradation. Ubiquitination at the single non-destabilizing site does not allow for proteasomal degradation. These results suggest that ubiquitin-mediated destabilization can reveal an obligate unstructured region in substrates that otherwise lack such a region. Furthermore, ubiquitination at sensitive sites results in more rapid degradation of these barstar variants when a proteasome-engageable unstructured tail is fused to their C termini (21).

This previous work clearly demonstrates that ubiquitin-mediated changes to the protein landscape can play an important role in proteasomal selectivity and processing; it did not, however, uncover the molecular mechanisms through which these site-specific effects arise. Here, we interrogate the molecular mechanisms of ubiquitin-induced changes for these same single-lysine variants of barstar. We investigate differences in the intrinsic dynamics of these regions within barstar and differences in how the protein responds to ubiquitination at these individual sites. We employed two sets of complementary approaches: 1) NMR and hydrogen-deuterium exchange mass spectrometry to characterize the equilibrium conformational fluctuations of the substrate protein in the presence and absence of ubiquitin, and 2) molecular dynamics (MD) simulations to track the position of every atom in a barstar, in the presence and absence of ubiquitin, starting from its native conformation over the timescale of microseconds.

We find that ubiquitination has only subtle effects on the native structure of barstar. Ubiquitination at the sensitive sites, however, selectively increases fluctuations that expose barstar’s C terminus. While both of the sensitive sites arise in regions of barstar with relatively high conformational flexibility, the observed destabilization appears to occur through different thermodynamic mechanisms. By contrast, ubiquitination at the non-sensitive site has a protective effect on barstar’s C terminus. Thus, the effects of ubiquitination at each site are highly dependent on the local context. This mechanistic understanding of the site-specific effects of ubiquitination should aid in developing a predictive understanding of the energetic consequences of individual ubiquitination events, and also contribute to our understanding of the ways in which aberrant lysine targeting leads to disease (23–25).

## Results

### Mono-ubiquitination at all three sites does not alter the native conformation of barstar

As noted above, we previously characterized the energetic effects of ubiquitination at three distinct sites of the small protein barstar using single lysine variants in which all but one native lysine had been mutated to arginine. Ubiquitination at positions K2 and K60 (the sensitive sites) destabilizes barstar, while ubiquitination at K78 (the non-sensitive sites) has relatively little effect (Fig. 1A) (21); these effects appear sufficient to allow for proteasomal engagement and degradation. We used these same single lysine variants in this study.

**Figure 1:**
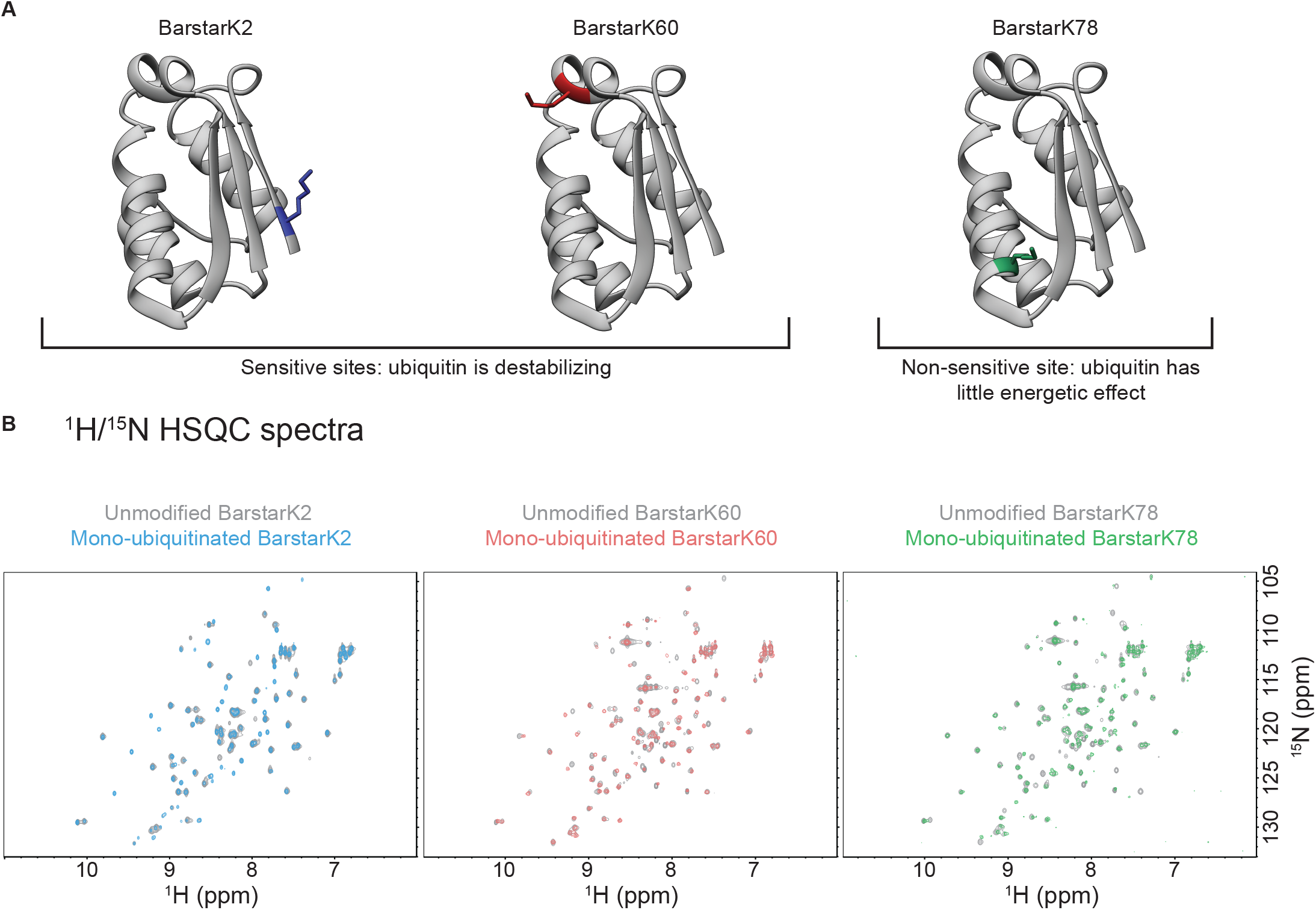
Overview of barstar sensitive and non-sensitive ubiquitination sites and NMR HSQCs of unmodified and mono-ubiquitinated barstar variants. (**A**) Ribbon diagram (PDB ID: 1BTA) depicting the position and surface topology of barstar lysine 2, a ubiquitin-destabilized site located in a β-strand, barstar lysine 60, a ubiquitin-destabilized site located in an ɑ-helix, and barstar lysine 78, a site located in an ɑ-helix that does not experience substantial destabilization upon ubiquitination. **(B)** NMR ^1^H/^15^N HSQC spectra depict the chemical shifts of amides from each residue in unmodified (gray) and overlaid mono-ubiquitinated barstarK2 (blue), barstarK60 (red) and barstarK78 (green). The similarity between the spectra of the unmodified barstars and their mono-ubiquitinated counterparts indicates that mono-ubiquitination does not substantially alter the native conformation of barstar, even at the two sensitive sites.

To determine whether ubiquitination alters the conformation of barstar, we first turned to NMR spectroscopy. HSQC NMR spectra were obtained for all three unmodified variants: barstarK2, barstarK60, and barstarK78 (Fig. 1B). All three show very similar spectra that closely resemble the published spectrum for wild-type barstar under slightly different conditions. Using the previously determined wild-type assignments (26, 27) and HNCA and HNCACB triple-resonance experiments, we assigned 70/88 barstar amides in unmodified barstarK60 to distinct peaks in the HSQC spectrum (BMRB ID: 50539). These assignments were easily transferable to the other two variants.

We next examined the HSQC spectra for the ubiquitin-modified variants. ^15^N-labeled barstar variants were ubiquitinated with unlabeled, fully methylated ubiquitin (which increases yield of mono-ubiquitination and has been shown to induce the same energetic effects on barstar as non-methylated ubiquitin (21)). HSQC spectra reveal well-dispersed peaks characteristic of well-folded proteins (Fig. 1B). Ubiquitin-induced perturbations to the conformation, as measured by the number and extent of chemical shift changes, appear to be relatively small for all three variants. These subtle effects are consistent with previous observations that, despite their destabilization and dramatic effects on proteasomal degradation, mono-ubiquitinated barstarK2 and barstarK60 unfold cooperatively and can still bind barstar’s binding partner, barnase (21). For barstarK2 and barstarK60, the only notable changes observed in the HSQC are largely limited to residues directly surrounding the site of the modification and near the C terminus. By contrast, mono-ubiquitinated barstarK78 exhibits fewer notable peak shifts near the site of ubiquitin modification.

In sum, these data indicate that all barstar variants adopt the native barstar-like fold in both their unmodified and mono-ubiquitinated states. In addition, an examination of barstar’s structure (PDB 1BTA) reveals no obvious basis for the observed differences in stability across the three variants. For instance, local contact density in the vicinity of a given ubiquitination site does not correlate with the sensitivity of that site to ubiquitination (Fig. S1). The observed energetic and functional changes to barstarK2 and barstarK60 must therefore arise from shifts in the population of high-energy, partially or globally unfolded states on the landscape.

### Conformational flexibility in the barstar native state

To explore the mechanistic basis for the energetic differences observed upon ubiquitination, we first set out to characterize the conformational dynamics of unmodified barstar using complementary computational and experimental approaches. Do the regions containing sensitive sites (the K2 strand and K60 helix) differ in their conformational dynamics from the non-sensitive site (the K78 helix)?

We performed extensive all-atom molecular dynamics simulations of unmodified barstar, starting from the conformation observed in the NMR structure, PDB entry 1BTA (27) (~30 μs in aggregate; see Materials and Methods). As expected, regions corresponding to portions of the sequence with secondary structure (α helices: residues 15–25, 34–43 and 68–80; and a β strand, residues 50–54) exhibit smaller backbone fluctuations compared to regions of the sequence lacking secondary structure (residues 7–14 and 25–33) (see Materials and Methods; Fig. 2A).

**Figure 2:**
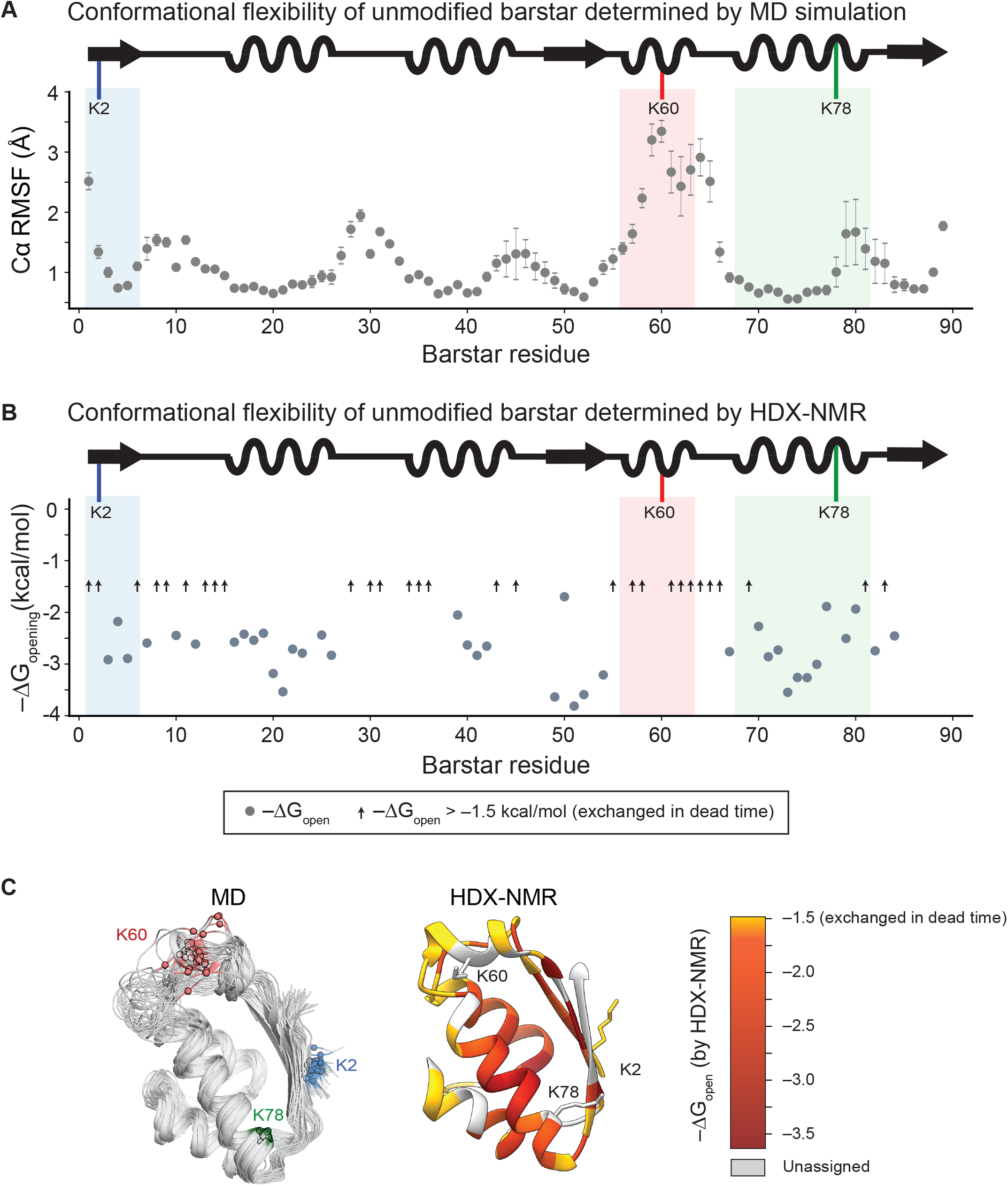
Barstar sensitive sites exist within regions of high intrinsic conformational flexibility in the unmodified protein. **(A)** Sites sensitive to ubiquitination occur within relatively flexible regions of barstar, as measured by the root-mean-square fluctuation (Å) of each Cα atom about its average position across six independent, 5.0-μs simulations of unmodified, wild-type barstar. Strand 1 (containing K2) is shaded in blue, helix 3 (containing K60) is shaded in red, and helix 4 (containing K78) is shaded in green. Error bars represent the SEM (*n* = 6). **(B)** Free energies of opening (ΔG_opening_) (gray circles) calculated from protection factors obtained by HDX-NMR of unmodified barstarK60 and plotted by barstar residue number (*n* = 1). Amide residues that exchange faster than the 9.2-minute dead time of the experiment are depicted by black arrows. Again, strand 1 (containing K2) is shaded in blue, helix 3 (containing K60) is shaded in red, and helix 4 (containing K78) is shaded in green. **(C)** Sites of backbone flexibility in barstar, as measured through MD and HDX-NMR. Left: overlapping simulation snapshots sampled every 100 ns after removing the first 1.0 μs (for equilibration) from a representative simulation of barstar. Spheres represent the Cɑ atom of each ubiquitination site. Right: ΔG_opening_ values from HDX-NMR mapped onto the NMR structure of barstar (PDB ID: 1BTA).

Two exceptions to this general trend arise—in helix 3 (residues 55–63), which rapidly loses its initial helicity and adopts many loop-like conformations, and in strand 1 (residues 1–5), which occasionally unbinds from its neighboring strand. Notably, the sensitive sites (K60 and K2) map to these regions, while the non-sensitive site (K78) does not. Throughout the course of each simulation, K2 and K60 form fewer contacts with neighboring residues compared to K78 (Fig. S1). Thus, these simulations indicate that the two sensitive sites occur in regions of greater conformational flexibility compared to the non-sensitive site.

Experimentally, we evaluated the fluctuations in unmodified barstar using hydrogen-deuterium exchange monitored by NMR (HDX-NMR). ^15^N-labeled barstarK60 was diluted into D_2_O and HSQC spectra were taken over a period of ~22 hours. Using our amide backbone assignments, we followed the decrease in the individual amide proton intensity (peak volume), and calculated the observed exchange rate for a given amide (*k*_obs_) (see Materials and Methods). From these rates, we then determined a protection factor, PF, using the known sequence-dependent exchange rates derived from unstructured peptides, *k*_*rc*_, (28–30), where PF = *k*_*rc*_/*k*_*obs*_. PF = 1 corresponds to a proton that exchanges at a rate expected for an unfolded amide, and PF > 1 corresponds to a protected amide. Given these limitations, we successfully determined rates of exchange and the corresponding PFs for 38 of the 88 amide sites (Fig. 2B). In agreement with our simulations, most regions of secondary structure contained well-protected residues. Again, the primary exception is helix 3, which exhibited a notable lack of well-protected amides compared to the other helices in barstar.

Together, these simulations and experiments suggest that intrinsic dynamics, or flexibility, may be an important feature governing the effects of ubiquitination events at individual sites within a protein (Fig. 2C). They raise the possibility that sensitive sites may tend to occur within regions of high intrinsic flexibility, while non-sensitive sites may occur in regions of lower flexibility (see Discussion).

### Ubiquitination at sensitive sites alters equilibrium fluctuations

To characterize the changes in barstar equilibrium fluctuations upon ubiquitination, we again used hydrogen-deuterium exchange, this time monitored by mass spectrometry (HDX-MS), following exchange at the peptide level (see Materials and Methods). Despite providing lower resolution than HDX-NMR, HDX-MS yields a structural picture of the changes in dynamics and requires smaller amounts of sample such that the experiment is feasible with our mono-ubiquitinated barstar variants. We performed these experiments for all three unmodified and mono-ubiquitinated barstar variants and analyzed peptides from both barstar and ubiquitin. In these experiments, exchange was monitored over a time course of four to eight hours. Fig. 3 summarizes the results.

**Figure 3:**
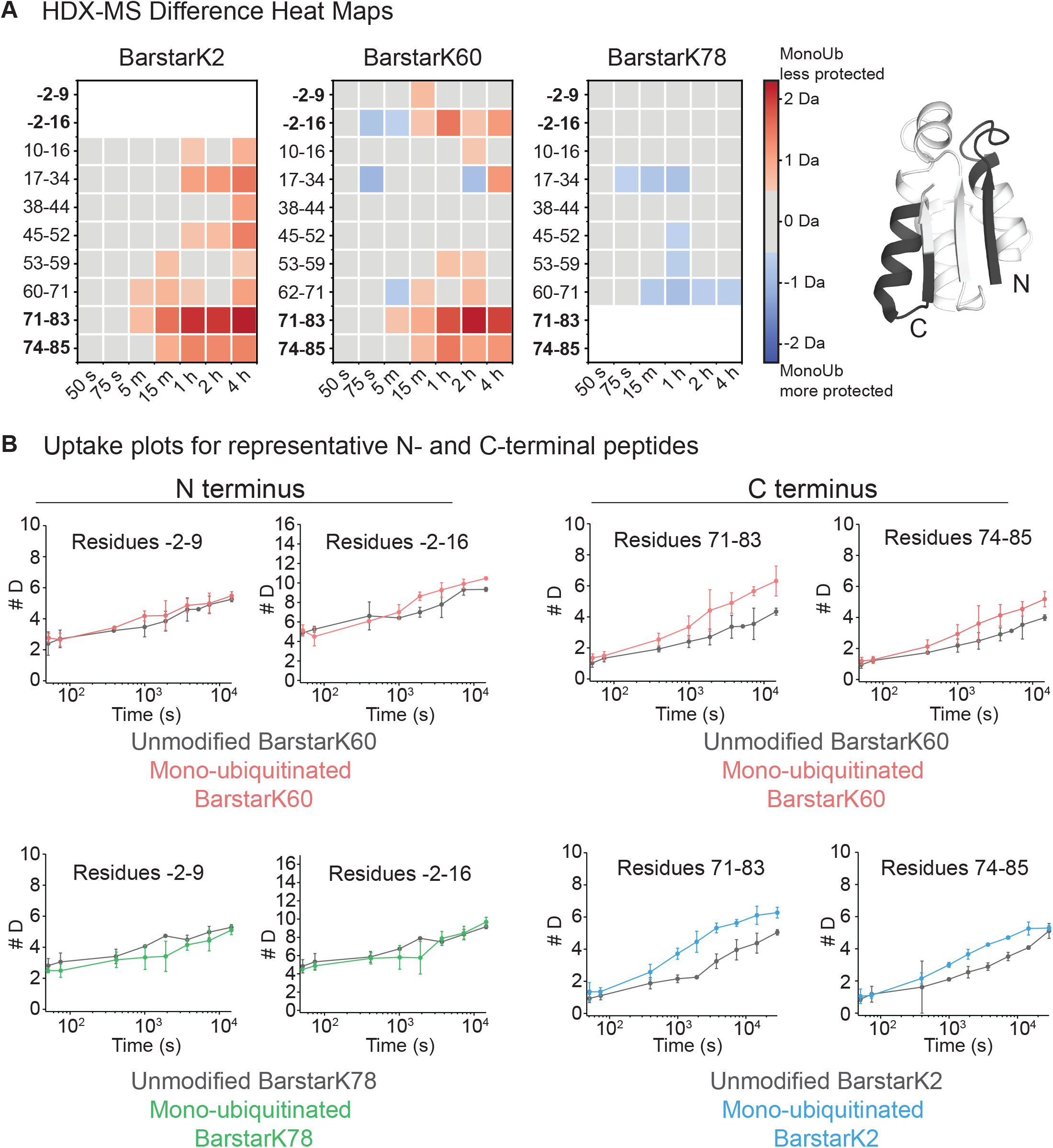
Ubiquitination at sensitive sites on barstar destabilizes its C terminus. **(A)** Heat maps from representative peptides depicting differences in deuteration between unmodified and mono-ubiquitinated barstarK2, barstarK60, and barstarK78 (*n* = 3 for both unmodified and mono-ubiquitinated samples), from HDX-MS experiments. Red indicates increased deuterium exchange in the mono-ubiquitinated compared to the unmodified state and blue indicates decreased deuterium uptake in the mono-ubiquitinated compared to unmodified state, both reported in Da. We consider a deuteration difference of at least ±0.5 Da between the unmodified and mono-ubiquitinated species meaningful (84) and use gray squares to indicate points falling below this 0.5 Da limit. The barstar C terminus, but not its N terminus, becomes significantly destabilized upon ubiquitination at sensitive sites. Peptides containing the site of the modification could not be analyzed because the ubiquitin ‘scar’ prevented accurate mass identification. Heat maps for the full peptide data sets are available in **Fig. S2. (B)** Deuterium uptake plots not corrected for back exchange from representative peptides from the N- and C-terminal regions of unmodified (grey) and mono-ubiquitinated (blue) barstarK2 (*n* = 3 for each protein state), unmodified (grey) and mono-ubiquitinated (red) barstarK60 (*n* = 3 for each protein state), and unmodified (grey) and mono-ubiquitinated (green) barstarK78 (*n* = 3 for each protein state). Error bars represent the standard deviation of replicates, and points are plotted at the average exchange time. Complete HDX-MS dataset is available as **source data**.

For all three variants, the majority of barstar peptides showed similar behavior in their ubiquitinated compared to unmodified states (Fig. 3A, Fig. S2). Only a small number of peptides exhibit significant differences in deuterium uptake between unmodified and mono-ubiquitinated barstar (Fig. 3B, Fig. S3). Both mono-ubiquitinated barstarK2 and barstarK60 show an increase in exchange compared to their unmodified counterparts that localizes to the C terminus and thus appears to rely on an allosteric network. This increased exchange, observed across multiple distinct C-terminal peptides in both barstarK2 and barstarK60, indicates a lower local stability upon ubiquitination. We could not analyze these same peptides for mono-ubiquitinated barstarK78 (the non-sensitive site) because the proteolysis scar (i.e. any ubiquitin residues still attached to the isopeptide bond) remaining from the ubiquitin modification prevented accurate peptide mass analysis/identification. At the N-terminus, we do not observe any notable changes for barstarK60 and K78, which show nearly identical exchange profiles between unmodified and mono-ubiquitinated samples (Fig. 3B). Again, due to the ubiquitin scar, we could not analyze N-terminal peptides for mono-ubiquitinated barstarK2.

These changes at the C terminus are particularly intriguing because such flexibility may be responsible for the observed proteasomal processing of these destabilizing variants. For small proteasome substrates, the proteasome frequently engages at either terminus (12). Our previous biochemical evidence implicates the C terminus as the probable site of degradation initiation for barstarK2 (21).

Monitoring hydrogen exchange by mass spectrometry allows us to also follow peptides from ubiquitin, both in isolation and attached to barstar (See Materials and Methods). Interestingly, these peptides show no discernible change in extent or time course of exchange over the eight-hour experiment, indicating that any changes to ubiquitin are not detectable in this time window (Fig. S4). This is surprising because, due to thermodynamic coupling, destabilization should be reciprocal. Nonetheless, ubiquitin possesses unusually high thermodynamic stability (31), and this high stability may serve to protect ubiquitin from unfolding and losing its signaling integrity upon conjugation to its myriad of targets in vivo (see Discussion).

Taken together, our HDX-MS and NMR analyses indicate that, while the three barstar variants all adopt the same native conformation, ubiquitination affects the conformational dynamics of each barstar variant differently. Specifically, the primary effects of ubiquitination at sensitive sites K2 and K60 occur in the C terminus, which experiences increased fluctuation upon ubiquitination. These results indicate that the effects of ubiquitination at these two sites propagate allosterically. The observed destabilization thus arises from these changes in native-state dynamics rather than changes to the native structure, consistent with our previous studies that showed slow proteasomal degradation rates suggesting a required conformational fluctuation.

These HDX-MS experiments raise further mechanistic questions: if ubiquitin-induced destabilization primarily affects the flexibility and/or exposure of the substrate C terminus, how do modifications at sites far from the C terminus induce these effects? Conversely, how can a modification at a site within the C terminus, such as the non-destabilizing K78 modification, not destabilize the protein?

### Thermodynamic basis for site-specific effects of ubiquitination

To probe the molecular mechanism by which ubiquitination can site-specifically modulate the barstar conformational landscape, we again turned to molecular dynamics simulations. For each barstar variant, we modeled an isopeptide bond to attach ubiquitin at the appropriate lysine site. Different potential lysine side-chain rotamers affect the orientation of ubiquitin with respect to barstar (Fig. S5A). For each variant we selected the rotamer that maximized the distance between the centers of mass of barstar and ubiquitin (Fig. S5B, see Materials and Methods). We then verified that simulations starting from these conformations could sample a wide variety of other conformations without becoming trapped in local minima, suggesting that the simulations are well-converged (Fig. S6). For all three variants, the ubiquitin structure typically moves quickly with respect to barstar (Fig. S6), and we only occasionally observe the formation of surface–surface interactions between the two proteins. All ubiquitinated variants sample fluctuations similar to those observed in simulations of unmodified barstar, with the secondary structure elements fluctuating less than regions composed of loops (Fig. S7).

The most notable difference in conformational fluctuations occurs in monoUb-barstarK60, where residues within helix 3 (residues 55-68) displace less from their average position (i.e. fluctuate less) (Fig. 4A, Fig. S7). A second difference occurs in the N terminus (residues 1–6) of monoUb-barstarK2, where residues 1–3 displace more from their average position (Fig. 4B, Fig. S7). Importantly, within helix 3, monoUb-barstarK2 and monoUb-barstarK78 behave more similarly to unmodified barstar than to monoUb-barstarK60, and at the N terminus, monoUb-barstarK60 and monoUb-barstarK78 behave more similarly to unmodified barstar than to monoUb-barstarK2. Thus, our results indicate that ubiquitination at either of the two sensitive sites affects conformational fluctuations local to those sites, while ubiquitination at the non-sensitive site has little effect on local flexibility. We next set out to identify the atomic-level changes in barstar that underlie these observed changes in flexibility.

**Figure 4:**
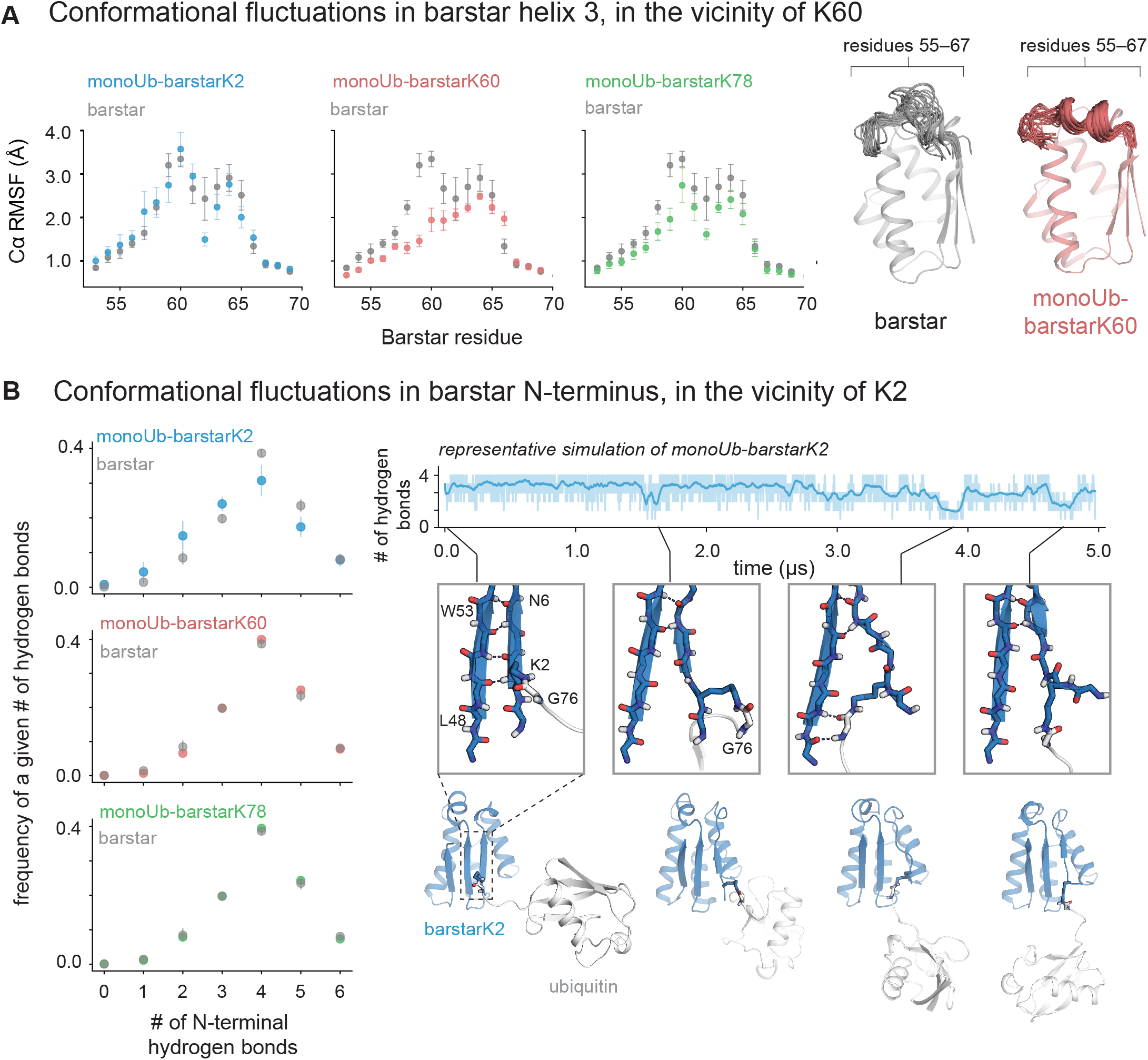
Distinct thermodynamic mechanisms of ubiquitin-mediated destabilization in barstarK60 and barstarK2. **(A)** Ubiquitination of K60 reduces local conformational heterogeneity in the vicinity of the modification. Simulations reveal substantially reduced backbone fluctuations about the average structure, as measured by the root-mean-square fluctuation (RMSF) of each Cɑ atom for the residues making up helix 3 (55–68). Error bars represent the SEM. (*n* = 6 simulations per condition). On right, simulation snapshots representing residues 55–68 of barstar sampled every 100 ns from a representative simulation are overlaid on the simulation starting structure. **(B)** Ubiquitination of K2 destabilizes interactions between the N-terminal β-strand of barstar and its neighboring β-strand (residues 50–53). For each simulation, we report the fraction of time a given number of hydrogen bonds (between 0 and 4) forms between residues 1–6 and 48–53. These histogram-like plots (left) reveal that ubiquitination of K2 *increases* the fraction of the time a simulation spends with a single hydrogen bond and *decreases* the fraction of the time a simulation spends with three hydrogen bonds. Trace from a representative simulation (top) shows the number of hydrogen bonds between barstar strand 1 and strand 2 over time in the simulation. Thick trace represents a sliding average of simulation values smoothed over a 50-ns window. Ribbon diagrams illustrate conformations from a representative simulation of ubiquitinated barstarK2 in which the hydrogen bonds between strand 1 and strand 2 are disrupted. In certain cases, ubiquitin’s flexible C-terminus occasionally forms intermittent non-native hydrogen bonds with barstar strand 2 when strand 1 is displaced. Ubiquitination at K2 therefore directly disrupts the three-strand β-sheet of barstar.

In simulations of monoUb-barstarK60, helix 3 samples a narrower range of conformations than it does in simulations of barstar alone or other monoubiquitinated variants. Specifically, its average simulated structure contains two full helical turns (Fig. S8), and it fluctuates less about its average structure than do the other barstar variants (Fig. 4A). By contrast, and as described above, unmodified barstar often spontaneously samples non-helical conformation, as does monoUb-barstarK2, and to a lesser extent, monoUb-barstarK78. Taken together, these results imply that ubiquitin limits the number of conformations helix 3 samples when attached to K60 but not when attached at other sites. Thus, the substantial reduction in number of conformations sampled introduces an entropic penalty that destabilizes the native structure of barstar.

In simulations of monoUb-barstarK2, strand 1 separates from its neighboring strand more often than it does in simulations of barstar alone or other monoubiquitinated variants (Fig. 4B). In particular, it adopts conformations in which strand 1 forms fewer backbone–backbone hydrogen bonds with strand 2, thereby disrupting the beta sheet of barstar. Intriguingly, strand 1 occasionally separates from strand 2 across simulations of all variants, with and without ubiquitin; these excursions persist for longer periods of time in monoUb-barstarK2 (Fig. S8). Moreover, a close examination of our simulations reveals that, for monoUb-barstarK2, the isopeptide linkage sometimes moves into the space between strand 1 and strand 2, thereby prolonging the amount of time monoUb-barstarK2 spends in this strand-displaced conformation. These observations are supported by our HDX-MS studies, which show that peptides encompassing strand 2 undergo increased exchange in the monoUb-barstarK2 background (Fig. 3A, Fig. S3). We therefore propose that ubiquitination at K2, but not at other sites, leads to direct disruption of the beta sheet in barstar, introducing an enthalpic penalty that destabilizes barstar.

In sum, these simulations indicate that while both sensitive sites arise in regions of local conformational flexibility, ubiquitination at each site induces destabilization through different molecular mechanisms—dominated by entropic effects (i.e. number of conformational states) in the case of K60 and by enthalpic effects (i.e. number of hydrogen bonds) in the case of K2. Importantly, the properties of the local conformational ensemble, rather than of the ground-state structure, appear to determine whether ubiquitin has an effect on each particular site. In each case, ubiquitination globally destabilizes the native state of barstar.

Notably, our simulations do not detect the concomitant exposure of the substrate C terminus observed in our HDX-MS experiments, likely because C-terminus exposure occurs on timescales associated with partial unfolding, which are beyond the scope of our simulations. Nevertheless, our data suggest that changes in the local flexibility near the two sensitive sites globally destabilize the native state of barstar, allosterically leading to increased exposure of the substrate C terminus.

How do these changes compare to the effects of ubiquitination at the non-sensitive site, K78? Across multiple independent simulations, mono-ubiquitinated barstarK78 samples a transient non-native hydrogen bonding network between the C-terminal β strand of barstar and the flexible C terminus of ubiquitin (Fig. 5A), contributing an additional strand to the core β sheet of barstar. Moreover, the strand contributed by ubiquitin occludes the barstar C terminus, substantially reducing its solvent exposure in simulation (Fig. S8C). These results provide an exciting molecular explanation for the lack of destabilization observed in this non-sensitive variant, and this increased stability at the C terminus may prevent barstarK78 from undergoing proteasomal degradation (21).

**Figure 5:**
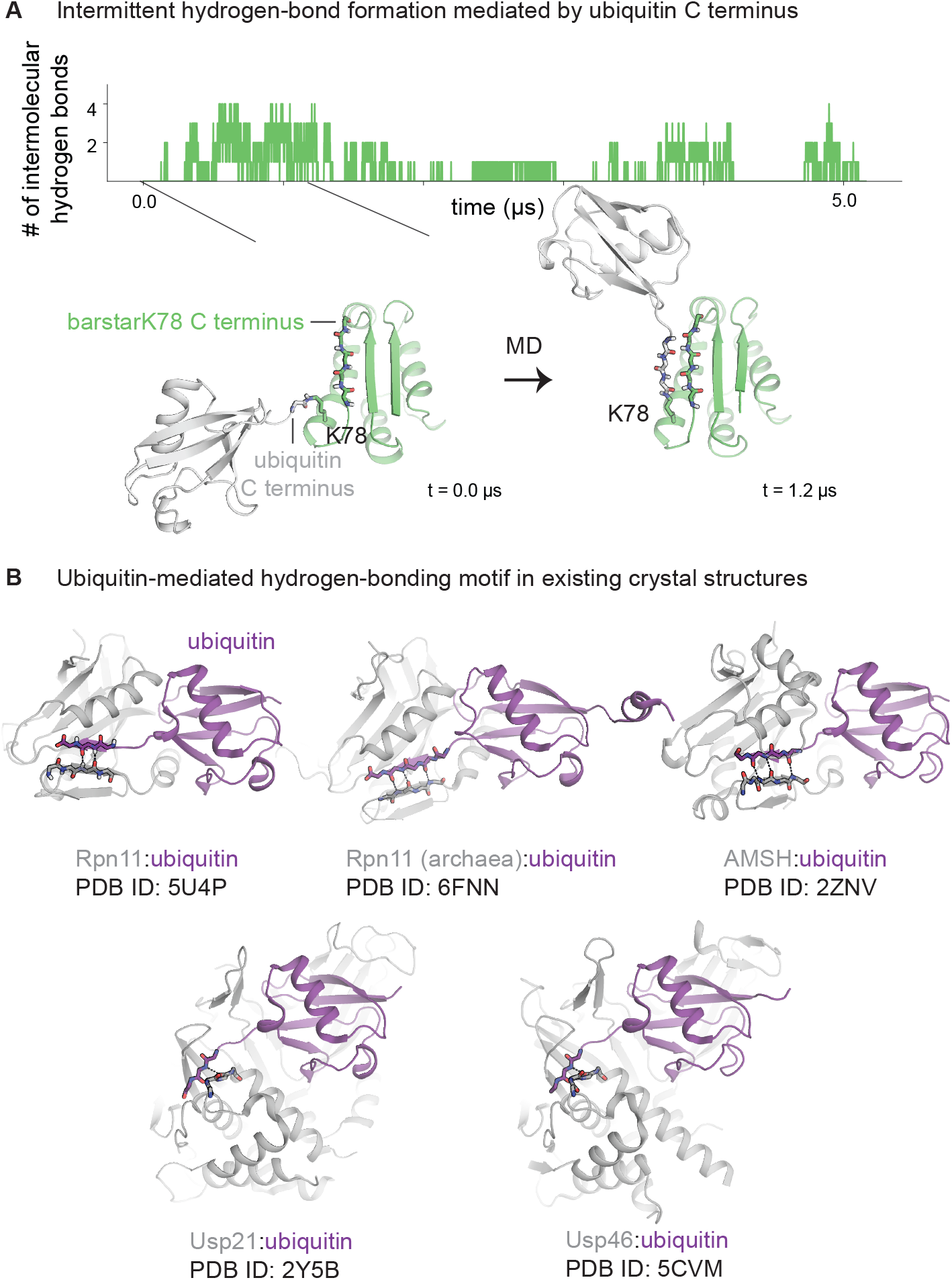
Non-native hydrogen bonding with ubiquitin’s C terminus protects barstarK78’s C terminus. **(A)** Ubiquitination at K78 protects the C-terminal β-strand of barstar, potentially stabilizing the substrate and protecting it from proteasomal targeting. Over the course of a simulation of barstar monoubiquitinated at K78, the ubiquitin C-terminus forms transient hydrogen bonds with the C-terminus of barstar, effectively adding a fourth β-strand to the existing β-sheet. These hydrogen bonds occur consistently across all six independent simulations of K78-ubiquitinated barstar. **(B)** The ubiquitin C-terminal tail constitutes a protein recognition and binding site. Crystal structures of other ubiquitin-bound protein complexes also contain the same interaction motif observed in simulations of K78-ubiquitinated barstar. These include several ubiquitin–deubiquitinase interactions.

This non-native hydrogen-bonding motif revealed in our simulations bears a striking resemblance to a motif observed in structures of a wide range of ubiquitin–protein binding interactions, including multiple classes of deubiquitinases (Fig. 5B). In all of these structures, the C-terminal tail of ubiquitin (typically residues 74–76) forms backbone–backbone hydrogen bonds with a β strand of the substrate protein (32–36). Thus, the isopeptide linkage, along with the local topology near the site of ubiquitination, facilitates specific interactions of ubiquitin with the substrate.

Aside from this non-native hydrogen-bonding motif, long-lived, non-covalent interactions between barstar and ubiquitin form only rarely in simulation (Fig. S6, Fig. S9). Moreover, neither barstar nor ubiquitin exhibit regions of increased protection from hydrogen exchange in HDX-MS experiments of ubiquitinated variants. Global analysis of our simulations shows that long-range barstar–ubiquitin interactions occur with a frequency of at most ~20% of the time (Fig. S9) and often involve the ubiquitin Ile44 surface hydrophobic patch which has been extensively characterized as a ubiquitin-protein binding motif (37). Intriguingly, in the case of monoUb-barstarK2, ubiquitin occasionally adopts an orientation that allows its C terminus to move between strands 1 and 2 of barstar (Fig. 4B), raising the possibility that a particular ubiquitin orientation might contribute to its destabilizing effect at this modification site. However, the consequences of these transient barstar–ubiquitin surface interactions for ubiquitin’s modulation of the barstar conformational landscape remains unclear from simulation alone.

Experimentally, we found that the ‘bulk’ of ubiquitin does in fact appear to play a role in energetic changes upon ubiquitination. We took advantage of the recently re-engineered deubiquitinase LbPro*, which collapses ubiquitin modifications to only the C-terminal GG residues of the proximal ubiquitin, leaving an isopeptide-linked GG scar (38). We used a previously established native-state proteolysis assay in which the observed rate of proteolysis by thermolysin is directly related to the free energy of partial unfolding (ΔG_proteolysis_) (21–39, 40). In this experiment, we find that LbPro*-treated monoUb-barstarK2 and monoUb-barstarK60 exhibit proteolysis kinetics intermediate between the mono-ubiquitinated and unmodified species (Fig. S10). Therefore, the LbPro*-treated species are destabilized to a lesser degree than the mono-ubiquitinated variants but are still slightly destabilized compared to the unmodified species. Thus, simply modifying the two sensitive-site lysines is not sufficient to propagate the allosteric effects of ubiquitination; the full ubiquitin modification (or at least more of the bulk) is necessary to induce the full magnitude of the observed energetic changes.

## Discussion

Using a combination of NMR, HDX-MS and MD simulation we interrogated the effects of site-specific ubiquitination on the conformational dynamics of barstar. Our results reveal site-specific mechanisms for ubiquitin-mediated changes. We find that these effects do not arise from changes in the ground-state structure of the substrate but rather are encoded by the substrate’s energy landscape. Moreover, these mechanistic changes appear to underlie observed differences in proteasomal degradation. We find that modifications at sensitive sites allow barstar to populate partially unfolded states that expose its C terminus. Ubiquitination at the non-sensitive site allows the barstar C terminus to sample a novel, and potentially stabilizing, interaction, as observed in simulation. The atomic-level changes associated with these effects arise from specific local properties of each modification site: for example, destabilizing effects can be dominated by either enthalpic contributions (i.e. by perturbing a substrate hydrogen bonding network) or entropic contributions (i.e. by reducing the number of configurations adopted by a region of the substrate).

We propose that, instead of dramatically changing the structure of the substrate, ubiquitin influences sub-global fluctuations on the energy landscape. This mechanism avoids the high energetic cost and potentially pathological consequences of fully unfolding a protein destined for degradation prior to its engagement with the 26S proteasome. These ubiquitin-induced modulations of the landscape allow the substrate to populate partially unfolded, high-energy conformation(s) sufficient for proteasomal engagement (21).

Our results suggest that ubiquitin induces fluctuations in the landscape that are either very sparsely populated or otherwise not accessible to the unmodified protein, thereby avoiding the population of substantially unfolded states that could template protein aggregation in cells. Thus, site-specific ubiquitin modulation might avoid aberrant activation of the protein quality control machinery or causing disease (41–44). In fact, previous studies have shown that the proteasome struggles to degrade aggregated proteins (45–47).

Due to thermodynamic coupling, barstar should reciprocally affect the energetics of ubiquitin. However, in our HDX-MS studies, we do not observe obvious barstar-induced effects on the landscape of ubiquitin. Our experiments only probe fluctuations within ~5 kcal/mol of the native state and, therefore, given the high stability of ubiquitin (~12 kcal/mol) (31), do not capture ubiquitin’s global and large subglobal fluctuations. Ubiquitin’s high stability may play an important role in providing tolerance and resistance to such energetic effects. Notably, ubiquitin has unusually high sequence conservation across multiple domains of life (48), but deep mutational scanning of ubiquitin has revealed its high tolerance to mutation (49, 50). Taken together, these results suggest that the stability of ubiquitin allows it to accommodate a range of perturbations, from a single point mutation to covalent linkage with a similarly sized protein, without compromising its fold. We therefore propose that its unusually high stability may have evolved in order to protect ubiquitin from unfolding and thus from losing signaling recognition and integrity upon conjugation to its myriad targets in vivo.

Importantly, we demonstrate that the bulk of ubiquitin (i.e. residues other than just the isopeptide-linked C-terminal GG residues) contributes to its site-specific effects, but those effects do not appear to depend on the formation of *specific* protein–protein interaction surfaces. Ubiquitin has several surface motifs that mediate ubiquitin–protein binding interactions (51, 52). Although one of those ‘patches’ forms occasional interactions with barstar in simulations of all three mono-ubiquitinated variants, neither experiments nor simulations identified surfaces of barstar or ubiquitin that became substantially protected in each other’s presence. Nevertheless, native-state proteolysis experiments with LbPro*-treated barstarK2 and barstarK60 revealed that removing the bulk of ubiquitin partially ablated the destabilizing effect of ubiquitination. This observation is consistent with an ‘entropic pulling’ effect, whereby transient interactions between the stably folded ubiquitin modification and substrate protein create a free energy gradient that generates a destabilizing net pulling force (53–55).

Critically, ubiquitin’s site-specific effects depend on how its covalent attachment perturbs the local dynamics of the substrate surrounding a given site. These observations provide an explanation for how ubiquitination at many distinct sites, on a wide variety of substrates, can encode the same cellular fates. At each of the two sensitive sites, ubiquitin acts via distinct mechanisms to alter the flexibility of barstar and thereby reduce the population of the native conformation with a concomitant increase in the population of a proteasome-engageable partially unfolded state. At the non-sensitive site, ubiquitin’s lack of effect correlates with the ability of its flexible C terminus to hydrogen bond with the barstar C terminus, contributing an additional strand to the barstar β sheet. Other substrates possessing a properly positioned, exposed set of backbone atoms might similarly interact with ubiquitin, also yielding protective effects. Interestingly, co-crystal structures of ubiquitin and various binding partners reveal similar structural motifs involving ubiquitin’s C terminus. Thus, the many conformational degrees of freedom of the native isopeptide bond, together with ubiquitin’s flexible C-terminal tail, confer substrate and site specificity, and studies of ubiquitination that employ non-native linkages may not capture such effects.

By extension, our results suggest that efforts to predict the effects of ubiquitination at a particular site need to account for the conformational flexibility of the substrate. Previous computational studies have sought to link structural properties of a ubiquitination site to ubiquitin’s eventual effect. For example, a bioinformatics analysis revealed that, unlike phosphorylation sites, which primarily arise in disordered regions, ubiquitination sites can arise in both ordered and disordered regions (16). A subsequent study employing coarse-grained simulations found that *high* contact density in the vicinity of the ubiquitin modification negatively correlates with thermodynamic stability (19), which differs from our own observation that the two sensitive sites on barstar arise in regions of *greater* intrinsic flexibility.

Our own preliminary bioinformatic analysis using a mass spectrometry screen for ubiquitinated substrates (56) (Fig. S11) reveals that degradative and non-degradative ubiquitination sites arising in structured regions of proteins are not readily distinguishable by the simple structural properties of the lysine residue, such as its secondary structure, number of neighboring contacts, or its depth from the surface. Our experimental and computational findings agree with and expand upon the results of these and other studies (20, 57)—the effects of ubiquitination depend strongly on local fluctuations in the vicinity of the modification site and cannot be inferred from the static structure alone. In addition, other cellular unfoldases that also recognize ubiquitinated proteins, such as p97, may respond to ubiquitin-induced energetic changes with additional consequences for proteostasis. Developing the ability to predict the effect of ubiquitination at a given site on any given substrate will have critical implications for the design of therapeutics via targeted ubiquitination, such as PROTACs (58).

Although we have used several different biophysical approaches to interrogate the effects of ubiquitination in atomic-level detail, our study comes with several caveats. In our simulations, we have not captured the entire process by which local destabilization near the site of ubiquitination allosterically affects the substrate C terminus, enabling substrate unfolding. We expect those changes to take place on timescales longer than those sampled in our unbiased simulations. Additionally, we have only examined the effects of ubiquitination on a single protein substrate, barstar, and further work is required to expand our results to other proteins, particularly those with physiological degradative and non-degradative ubiquitination sites. Finally, we have only examined the mechanistic effects of monoubiquitination, and characterization of the effects of ubiquitin chains with diverse lengths, linkages, and topologies is needed to fully understand the energetic effects conferred by ubiquitination in vivo (59).

Taken together, our work supports the idea that site-specific ubiquitination events induce distinct and consequential mechanisms for modulating a protein’s energy landscape. This site specificity allows a single post-translational modification to have a broad range of effects in cells, a phenomenon also associated with phosphorylation and glycosylation (16–60, 61). Site specificity of ubiquitin attachment is a built-in feature of the enzymatic conjugation and deubiquitination machinery in vivo, and the distinct mechanisms of ubiquitin-induced energetic effects represent a new layer of protein quality control and signaling in cells.

## Materials and Methods

### Purification of unmodified barstar variants

Purified mono-ubiquitinated barstarK2, barstarK60, and barstarK78 were prepared as described in (21). Only barstar variants used in LbPro*-treated native-state proteolysis experiments were fluorescein-labeled.

### Purification of ubiquitination enzymes

Ubiquitination enzymes *M. musculus* E1, *S. cerevisiae* Ubc4 E2, and *S. cerevisiae* Rsp5 E3 were purified as described in (21).

### Purification of ubiquitinated barstar variants

Purified mono-ubiquitinated barstarK2, barstarK60, and barstarK78 were prepared using methylated ubiquitin as described in (21) except that barstar variants were not fluorescein-labeled (with the exception of those prepared for LbPro*-treated native-state proteolysis experiments).

### Purification of ^15^N/^13^C-labeled proteins

*E. coli* BL21 Rosetta 2 (DE3) cells were transformed with either pEC076 (barstarK2), pEC062 (barstarK60), or pEC059 (barstarK78). Cells were grown in M9 minimal media prepared with ^15^N-labeled ammounium chloride as the sole nitrogen source. To prepare double-labeled samples, ^13^C glucose was also included as the main carbon source. Cells were grown to 0.4 < OD_600_ < 0.8 and induced with 1 mM IPTG overnight at 18°C. Bacteria were pelleted and resuspended in 50 mM HEPES pH 7.0, 150 mM NaCl, 0.5 mM TCEP supplemented with 1X Halt™ protease inhibitor cocktail (Thermo) and benzonase (Novagen). Resuspended cells were lysed by sonication and the lysate was clarified by centrifugation at 20,000 rcf, 4°C, 30 minutes. The substrate was first purified by Ni^2+^-NTA affinity chromatography using its N-terminal His_6_ tag. Clarified lysate was allowed to batch bind to HisPur™ Ni^2+^-NTA resin (Thermo), washed with 50 mM HEPES pH 7.0, 150 mM NaCl, 25 mM imidazole, 0.5 mM TCEP and eluted with 50 mM HEPES pH 7.0, 150 mM NaCl, 500 mM imidazole, 0.5 mM TCEP. Eluate was then run over an S200 16/60 size exclusion column (GE) pre-equilibrated with 50 mM HEPES pH 8.0, 150 mM NaCl, and 5 mM MgCl_2_. The peak corresponding to the full length His-MBP substrate was collected, and quantified by UV-Vis absorption at 280 nm before addition of 10% glycerol and flash freezing to store at −80°C for future use.

Before NMR data acquisition, samples were thawed, cleaved with HRV3C protease overnight at 4°C, and then run over an S75 16/60 size exclusion column (GE) equilibrated with 55 mM sodium phosphate pH 6.5 and 55 mM NaCl to collect the pure unmodified barstar peak. Samples were concentrated to 250 μL for data collection in a Shigemi tube. Typical final sample concentrations were in the range of 0.75-2 mM (unmodified barstar) and 75-200 μM (mono-ubiquitinated barstar). Pure, homogeneous, mono-ubiquitinated ^15^N-labeled barstar samples were purified using methylated ubiquitin as described in section 2.8.4. Samples were diluted with 10% D_2_O before NMR data acquisition.

### ^15^N/^1^H HSQC and ^13^C/^15^N/^1^H HNCA and HNCACB NMR data acquisition and processing

NMR experiments were recorded on a Bruker Inc. Avance II NMR spectrometer operating at 900 MHz and equipped with a CPTCI cryoprobe. The sample temperature was set to 298 K.

2D ^1^H-^15^N correlation spectra were recorded using the SOFAST HMQC method (62) using Bruker pulse sequence sfhmqcf3gpph. A total of 2048 pt and 256 pts were acquired in the ^1^H and ^15^N dimensions, respectively, with the ^1^H offset centered at 8.3 ppm and the ^15^N carrier frequency set to 120 ppm. The spectral widths were 16 ppm (^1^H) and 40 ppm (^15^N). The number of scans ranged from 4 to 32 depending on sample concentration, and the recycle delay was set to 0.5 sec.

3D HNCA and HNCACB spectra were recorded using the T1 optimized BEST-HNCA (63, 64) and BEST-HNCACB (64) methods with Bruker pulse sequences b_hncagp3d and bhncacbgp3d. For each experiment, a total of 1024 (^1^H), 64 (^15^N), and 256 (^13^C) pts were recorded with spectral widths of 14 ppm for ^1^H and 35 ppm for ^15^N. The ^13^C spectral width was set to 40 ppm for the HNCA spectrum and 80 ppm for the HNCACB spectrum. A ^1^H offset of 8.3 ppm was used, along with ^15^N and ^13^C carrier frequencies of 117.5, and 54 ppm, respectively. Four scans were signal averaged for each block, and the recycle delay was set to 0.5 sec. Each spectrum was recorded in approximately 12 hours.

The 2D and 3D data were processed with the NMRPipe software package (65) and analyzed using CARA (66).

### ^15^N/^1^H HSQC NMR hydrogen-deuterium exchange

15N-labeled unmodified barstarK60 was run over an S75 10/300 size exclusion column (GE) equilibrated with 50 mM sodium phosphate pH 6.5 and 50 mM NaCl. The pure unmodified barstarK60 peak was collected and concentrated to 250 μL for eventual data collection in a Shigemi tube. The sample was then lyophilized overnight. The sample was resuspended in equivolume D_2_O, and NMR data collection began immediately after re-dissolving the sample.

Amide proton exchange rates for unmodified barstarK60 were obtained by measuring intensities of peaks in a series of 2D ^1^H-^15^N correlation spectra recorded using the SOFAST-HMQC (62) method. For each spectrum, eight scans were signal averaged per block, resulting in an acquisition time of 20 mins. Sixteen spectra were recorded with initial time points of 9.2, 30.1, 50.9, 71.7, 92.5, 113,3, 134,1, 154.9, 175.7, 196.5, 217.3, 255.6, 516.4, 777.2, 1037.9, 1298.8 minutes after dissolution of lyophilized protein into D_2_O.

### Hydrogen-deuterium exchange mass spectrometry experiments

To prepare deuterated buffer, 5 mL of a 50 mM HEPES pH 7.0, 50 mM NaCl, 50 mM KCl, 10 mM MgCl_2_ buffer in a 50 mL conical flask was flash frozen in liquid nitrogen and lyophilized overnight. The lyophilized buffer was then resuspended in equivolume D_2_O and allowed to exchange for 6 hours at room temperature before flash freezing and lyophilization. This process was repeated for a total of three D_2_O resuspension and lyophilization steps. After the final lyophilization step, the lyophilized buffer was parafilmed and stored at −80°C for future use.

All three unmodified and all three mono-ubiquitinated barstar variants were purified via size-exclusion chromatography as described above and then diluted with 50 mM HEPES pH 7.0, 50 mM NaCl, 50 mM KCl, 10 mM MgCl_2_ buffer to a final concentration of 10 μM in preparation for mass spectrometry experiments described below.

All hydrogen-deuterium exchange mass spectrometry (HDX-MS) experiments were performed using a liquid handling robot (LEAP Technologies) connected to a Q Exactive mass spectrometer (Thermo). The liquid handling robot was programmed to initiate amide proton exchange at 20° C by diluting barstar or mono-ubiquitinated barstar to 1 μM into deuterated buffer (50 mM HEPES pH 7, 50 mM NaCl, 50 mM, KCl, 10 mM MgCl2) and then quenching exchange at various timepoints by adding low pH buffer (6 M urea, 200 mM Arginine, 100 mM TCEP, pH 2.5) and cooling to 1° C. Quench buffer was added to the exchanging sample in a 1:1 ratio by volume such that the final protein concentration = 0.5 μM. The liquid handling robot was programmed to collect time points at 30, 60, 300, 900, 1800, 3600, 7200, 14400, and 28800 seconds; however, actual quench times are reported as source data. Average quench times for each programmed time point are used in Fig. 3B. After quenching, the samples were directly subjected to an in-line proteolysis step using one pepsin-packed column (to generate barstar peptides) or one pepsin followed by one fungal protease-packed column (to generate ubiquitin peptides). Proteolysis was followed directly by liquid chromatography using a C4 trap column followed by a C8 analytical column eluted with a 10-100% acetonitrile gradient and identified via mass spectrometry. Peptide lists were generated from an MS/MS run performed on each replicate using either Proteome Discoverer (Thermo) or Byonic (Protein Metrics). For peptides from methylated ubiquitin, only peptides containing the di-methylation modification were analyzed. Peptide deuteration states and isoptopic distributions were then determined using HD Examiner 2 (Sierra Analytics) with manual adjustment to the HD Examiner peak identifications as needed. Data are reported as absolute mass increases (comparing unmodified and mono-ubiquitinated variants to one another within the same experiment) and are not corrected for back exchange.

To construct difference heat maps, absolute mass values for the unmodified variants were subtracted from absolute mass values for the modified variants at each time point. We performed the subtraction on experiments of the unmodified and modified variants taken on the same day, and then averaged the resulting difference values. We do not consider differences less than ±0.5 Da as meaningful; and those data points are colored gray in the heat maps. Similar approaches have been taken for analysis of HDX-MS data in the past (67). Additionally, we excluded peptides greater than 20 residues in length.

### Purification of LbPro*

The gene encoding LbPro* (38) was cloned into an expression vector with an N-terminal, HRV3C protease-cleavable His-MBP tag to create pEC129. *E. coli* BL21 (DE3) cells were transformed with pEC129, and grown in LB media to 0.4 < OD_600_ < 0.8 and induced with 1 mM IPTG at 37°C for 3 hours. Bacteria were pelleted and resuspended in 50 mM HEPES pH 7.5, 150 mM NaCl, 0.5 mM TCEP supplemented with 1X Halt™ protease inhibitor cocktail (Thermo) and benzonase (Novagen). Resuspended cells were lysed by sonication and the lysate was clarified by centrifugation at 20,000 rcf, 4°C, 30 minutes. The construct was purified by Ni^2+^-NTA affinity chromatography using its N-terminal His_6_ tag. Clarified lysate was allowed to batch bind to HisPur™ Ni^2+^-NTA resin (Thermo) washed with 50 mM HEPES pH 7.5, 150 mM NaCl, 25 mM imidazole, 0.5 mM TCEP and eluted with 50 mM HEPES pH 7.5, 150 mM NaCl, 500 mM imidazole, 0.5 mM TCEP. Eluate containing pure protein was then concentrated and quantified by UV-Vis absorption at 280 nm before addition of 10% glycerol and flash freezing to store at −80° C for future use. Retention of the N-terminal His-MBP tag did not affect enzymatic processing, as whole-protein mass spectrometry confirmed that deubiquitination reactions yielded a ubiquitin species lacking the C-terminal GG residues.

### LbPro* deubiquitination and native-state proteolysis experiments

Purified, mono-ubiquitinated samples prepared with methylated ubiquitin and labeled with fluorescein-maleimide on the single barstar cysteine at position 82 were treated with 20 uM LbPro* at 37°C for 1 hour in reaction buffer (25 mM HEPES pH 7.5, 150 mM KCl and 15 mM MgOAc) supplemented with 2 mM DTT. LbPro* treatment under these conditions generated a mixed population of mono-ubiquitinated and –GG-modified barstar. Samples were then batch bound with amylose resin (NEB) to remove the His-MBP-LbPro* enzyme, and the flow through was collected.

Samples were allowed to equilibrate back to room temperature overnight, and then native-state proteolysis experiments were performed with 0.2 mg/mL thermolysin protease (Sigma). Time points were taken a t = 0, 0:15, 0:30, 0:45, 1:00, 1:30, 2:00, 3:00, 5:00, 7:00 and 10:00 min and quenched with EDTA. Each time point was then run on a NuPAGE Bis-Tris gel (Invitrogen) and band intensities quantified using ImageJ. Proteolysis rates are proportional to the free energy of partial unfolding to the protease-cleavable state (ΔG_proteolysis_) and have been previously measured for unmodified and mono-ubiquitinated barstarK2, barstarK60, and barstarK78 (21).

### System setup for molecular dynamics simulation

We simulated barstar under four conditions: on its own and with a single monomer of ubiquitin covalently attached, via an isopeptide linkage, to one of three sites on barstar (K2, K60 and K78). We performed six independent ~5.0-μs simulations for each construct: unmodified barstar, monoUb-barstarK2, monoUb-barstarK60, and monoUb-barstarK78. For each simulation, initial atom velocities were assigned randomly and independently.

We initiated simulations of barstar alone from the NMR structure of barstar, PDB entry 1BTA (27). To model each of the three mono-ubiquitinated variants studied experimentally in this paper, we employed the crystal structure of ubiquitin, PDB entry 1UBQ (68). In order to place ubiquitin in close proximity to a given lysine residue on barstar, in PyMOL, we aligned the lysine side chain to the ubiquitin-linked lysine side chain K48 in chain B of PDB entry 3NS8, and we aligned the ubiquitin C-terminal glycine of 1UBQ to the corresponding atoms in chain A of 3NS8. We exported these two molecules as a combined structure from PyMOL and loaded the resulting PDB into Maestro (Schrödinger, Inc.), where we used the Build tool to form a bond between the lysine side-chain nitrogen atom and the glycine carbonyl carbon. We then reduced the charge on the lysine to neutral and performed a local minimization on the lysine and glycine residues. Finally, we ensured that the torsion angle about the isopeptide bond matched a planar, *trans* configuration. The lysine side chain is highly flexible and can adopt ≥ 20 potential side-chain rotamers. To select a model of isopeptide-linked monoUb-barstar for simulation, we picked the lysine rotamer that maximized the distance between the centers of mass of ubiquitin and barstar. We prepared the resulting structures in Maestro with the Protein Preparation tool, retained titratable residues in their dominant protonation state at pH 7, and added neutral acetyl and methylamide groups to cap the free N- and C-termini, respectively, of the protein chains.

To place the prepared structures in a water box for simulation, we relied upon the LEaP program in AMBER (69). We solvated barstar using the LEaP command *solvateBox* with a buffer of 20 Å extending from the protein surface, resulting in a water box with initial dimensions of 75 Å × 75 Å × 75 Å. We next added sodium and chloride ions to the system to a concentration of 150 mM using the command *addIonsRand* with a minimal separation distance of 5 Å, for a total system size of 53,317 atoms. For the ubiquitinated barstar variants, we followed a similar protocol to the one outlined above, resulting in a water box with initial dimensions of 102 Å × 102 Å × 102 Å for each system. After adding ions, the total number of atoms was 134,218 for monoUb-barstarK2; 134,222 for monoUb-barstarK60; and 134,110 for monoUb-barstarK78. We also used Dabble (70) to guide generation of input files for LEaP.

For all simulations, we used the AMBER ff14SB for protein atoms, the TIP3P parameter sets for ions, and the TIP4P-D model for waters (71–73). We derived parameters for the isopeptide bond manually by explicitly assigning parameters for a standard peptide bond to specific atoms involving the side-chain residues of the modified lysine and glycine residues (after assigning unique atom types to those atoms), following a scheme described by (57). Parameter sets available with the source data for this paper.

### Simulation protocols

We performed our simulations using the Compute Unified Device Architecture (CUDA) version of Particle-Mesh Ewald Molecular Dynamics (PMEMD) in AMBER on single graphical processing units (GPU) (74). Simulations were performed using the AMBER18 software (69). Briefly, after three rounds of steepest descent minimization, followed by conjugate gradient minimization, systems were heated from 0 K to 100 K in the NVT ensemble over 12.5 ps and then from 100 K to 310 K in the NPT ensemble over 125 ps at 1 bar, with 10.0 kcal mol^−1^ Å^−2^ harmonic restraints applied to non-hydrogen protein atoms. Systems were then equilibrated at 310 K in the NPT ensemble at 1 bar, with harmonic restraints on non-hydrogen protein atoms tapered off by 1.0 kcal mol^−1^ Å^−2^ starting at 5.0 kcal mol^−1^ Å^−2^ in a stepwise fashion every 2 ns for 10 ns, and then by 0.1 kcal mol^−1^ Å^−2^ starting from 0.5 kcal mol^−1^ Å^−2^ for 2 ns in a stepwise fashion for 10 additional ns. For simulations of barstar, the same protocols were used, except that equilibration ended after the first 12 ns (the last equilibration step employed harmonic restraints of 0.5 kcal mol^−1^ Å^−2^ for 2 ns). Production simulations were carried out in the NPT ensemble at 310 K and 1 bar, using a Langevin thermostat for temperature coupling and a Monte Carlo barostat with isotropic control for pressure coupling. We employed a 4-fs time step with hydrogen mass repartitioning (75) and constrained bond lengths to hydrogen atoms using SHAKE (76). Non-bonded interactions were cut off at 10.0 Å. Long-range electrostatic interactions were computed using particle mesh Ewald (PME) with an Ewald coefficient of approximately 0.28 and a B-spline interpolation order of 4. The FFT grid size was chosen such that the width of a grid cell was approximately 1 Å. Trajectory snapshots were saved every 200 ps.

### Analysis protocols for simulation

The AmberTools19 CPPTRAJ package (69, 77) was used to center and reimage trajectories, and Visual Molecular Dynamics (VMD) was used to visualize and analyze simulations (78). All aggregate analysis, as well as individual simulation traces, use data analyzed every 1 ns for computational efficiency, unless indicated otherwise. In certain cases, we show time traces from simulation smoothed with a rectangular moving average of window size of 20 ns. Plots involving simulation data were visualized using the PyPlot package from Matplotlib (79).

For all analysis in the manuscript that required structural alignment, we aligned barstar on its Cα atoms in regions with ordered secondary structure (residues 2–6, 13–24, 34–42, 49–54, 57–62, 66–80, and 84–88). We computed the root-mean-square fluctuation (RMSF) for each simulation by first determining the average structure of barstar across all simulations within a given simulation condition (i.e. that is, for each of the four barstar constructs) after removing the first 100 ns of simulation to allow for complete system equilibration; we then calculated the mean-square fluctuation about each Cα along the barstar sequence, using the average structure as the reference. Error bars shown in figures containing RMSF plots refer to the standard error (the standard deviation of RMSF values for each simulation, divided by the square root of the number of simulations per condition—in this case, 6).

To quantify the presence of native hydrogen-bonding interactions involving the barstar N terminus in simulation, we calculated the distance between each amide proton on strand 1 (residues 1, 3 and 5) and each carbonyl oxygen on strand 2 (residues 48, 50, 52 and 54), and for the corresponding pairs of carbonyl oxygen acceptors on strand 1 and amide proton on strand 2. In our analysis, a pair forms a hydrogen bond if the distance between the acceptor atom and the hydrogen is ≤ 2.2 Å; we visually verified that this fast approach identified the same sets of hydrogen bonds as those identified using VMD’s geometric criteria of a 50° donor– acceptor angle and a ≤ 3.5 Å distance between the acceptor and donor atoms. Again, we removed the first 100 ns of each simulation to allow for complete system equilibration.

We performed a similar analysis to quantify the presence of non-native hydrogen-bonding interactions that formed between the C terminus of ubiquitin and the C terminus of barstarK78. Specifically, we examined the formation of hydrogen bonds between residues 74–76 on ubiquitin and 86–88 on barstar. We found that one non-native hydrogen bond formed at this interface in 32% of simulations, on average, for monoUb-barstarK78 and two non-native hydrogen bonds formed with a frequency of at 11%, on average. We occasionally observed that up to four non-native hydrogen bonds formed between the two C termini, as demonstrated in Fig. 5 and Figure S8.

To calculate the solvent accessible surface area of the barstar C terminus, we used the *sasa* command in VMD. We found the solvent accessible surface area of the atoms that make up barstar or the ubiquitin C terminus (residues 72–76) and used the *restrict* option to limit the solvent-accessible points considered to those contributed by the barstar C terminus (residues 83–89).

To generate intermolecular contact maps between ubiquitin and barstar, we used the GetContacts package (80) (https://getcontacts.github.io/index.html). To obtain a list of all van der Waals interactions, we ran the get_dynamic_contacts.py script on residues 1–89 of barstar, and residues 1–76 of ubiquitin, and then we obtained the frequency of those interactions by using the get_contact_frequencies.py script. Resulting heatmaps represent the frequency of the observed interactions. The brightest color on each color bar corresponds to an interaction frequency of 15% across all simulations in a given condition.

In the supplement, we also report the number of residue contacts near a given ubiquitinated lysine residue as the number of residues within 4 Å of the side chain of the lysine of interest, calculated using VMD. We performed this calculation on the static structure of barstar, as well as for simulated barstar. Error bars correspond to the standard error of the mean, where the mean corresponds to the average number of residue contacts across each of the six simulations of unmodified barstar.

### Analysis of ubiquitination sites identified by proteomics screen

Although we found that local properties of ubiquitination sites in the experimental structure of barstar (contact density and secondary structure) do not correlate in an obvious way with changes in stability due to ubiquitination, we asked whether such trends might exist for ubiquitination sites in other proteins. Indeed, an analysis on a manually curated dataset suggests that this might indeed be the case (19). To determine whether sites of ubiquitination, found within structured protein regions, differ in their local properties depending on whether they encode either degradative or regulatory effects, we performed a structural analysis of an existing proteomics dataset from (56).

Briefly, Gendron and colleagues used mass spectrometry to detect ubiquitin modifications (i.e., lysine residues modified by two glycines) across the proteome under different cellular conditions. Specifically, their study monitors the abundance of ubiquitination events upon inhibition of protein homeostasis. While the authors employ many different inhibitors of protein homeostasis in their study, here we used only data based on epoxomicin inhibition of the proteasome. Ubiquitination at a given site often determines whether a protein will undergo either degradative processing by the proteasome or regulation by some other means: degradative sites are considered to be ubiquitinated sites whose abundance increases upon proteasomal inhibition, and regulatory sites are ubiquitinated sties whose abundance decreases upon proteasomal inhibition.

We analyzed data found in Table S2 (and shown as a heatmap in Figure 2B) of ref. (56). These data represent the 200 most abundant sites for which spectral counts of dGG-containing peptides changed by more than 2-fold upon proteasomal inhibition by epoxomicin. We first categorized the sites as either ‘degradative’, as they increased in abundance upon inhibition, or ‘regulatory’, as they decreased in abundance upon inhibition. Uniprot IDs corresponding to proteins containing dGG sites were mapped to their entries in the Protein Data Bank using the Retrieve ID/mapping tool in Uniprot (https://www.uniprot.org/uploadlists/). For sites increasing in abundance with epoxomicin concentration, 46 of the 66 Uniprot IDs mapped to 895 PDB IDs. For sites decreasing in abundance with epoxomicin concentration, 59 of the 83 Uniprot IDs mapped to 1003 PDB IDs.

Subsequently, we employed the Biopython package (81) to extract the corresponding FASTA file associated with a particular Uniprot ID and to select the lysine residue within a PDB file that corresponded to the ubiquitin modification site specified in the dataset above. For each PDB file mapped to a particular Uniprot ID, we aligned each chain of the PDB file to the corresponding FASTA file for a particular Uniprot ID, using parameters for the PairwiseAligner() tool that minimize internal gaps. We then determined whether the site of ubiquitin modification existed within the sequence found in the PDB structure. Finally, for each site present within a PDB structure, we used the mkDSSP tool (https://slackbuilds.org/repository/14.1/academic/mkDSSP/; (82)) to classify its secondary structure as well as its relative accessible surface area (rASA). For each ubiquitination site, we selected one representative PDB file for more detailed structural analysis (the PDB whose site yielded the median rASA). This analysis yielded the depth of each residue side chain from the surface (http://mgltools.scripps.edu/packages/MSMS; (83)) as well as the number of residues within 4.5 Å of any non-hydrogen side-chain atom. Analysis scripts to be made available with source code.

In Fig. S11, the analyzed data represent 70 distinct ubiquitination sites detected through mass spectrometry; of these, 32 represent regulatory sites and 38 represent degradative sites; 1170 PDB structures were analyzed. Certain sites were represented by more than one PDB entry. In those cases, we selected the most common (i.e. the mode) secondary structure designation from DSSP.

## Author Contributions

E.C.C., N.R.L., and S.M. designed the research. E.C.C. performed the experiments. N.R.L. performed and analyzed molecular dynamics simulations. E.C.C. and N.R.L. analyzed and interpreted the combined experimental and computational datasets. J.M.L. assisted in HDX-MS data acquisition, optimization, and analysis and provided expertise in mass spectrometry. J.G.P. assisted in NMR data acquisition and analysis and provided expertise in NMR. B.C.M. assisted with protein purification and NMR data analysis. E.C.C., N.R.L. and S.M. wrote the manuscript. All authors edited and approved the manuscript.

## Acknowledgements

We thank all members of the Marqusee lab for helpful discussions and scientific guidance. We especially thank Helen Hobbs for assistance with HDX-MS data collection in the midst of a global pandemic. We thank Robin Betz for assistance with simulation preparation and Robert Best for advice on force-field parameters for simulation. We thank Andy Martin, Greg Bowman, and Meredith Rickard for helpful feedback. We also thank Ken Dong for helpful conversations about modeling the isopeptide bond; Jeff Morrow and Sierra Analytics for making HD Examiner licenses freely available for at-home use during shelter-in-place orders; and Berkeley Research Computing for providing computational resources and support through our use of the Savio cluster. This work was supported by the US National Institutes of Health: R01GM050945 to S.M. S.M. is a Chan Zuckerberg Biohub Investigator. N.R.L. is a UC Berkeley Miller Fellow. Funds for the 900 MHz NMR spectrometer were provided by the NIH through grant GM68933.

## Supplementary Information

**Figure S1:**
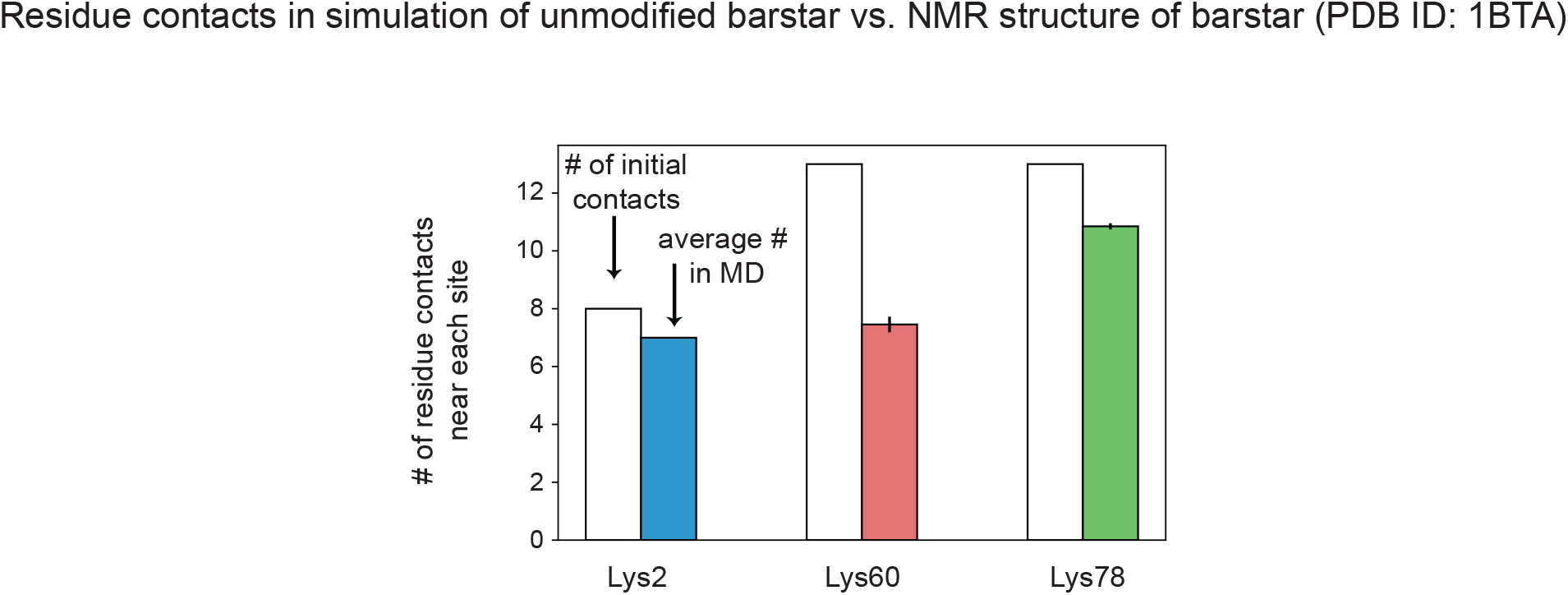
Native contact density at barstar ubiquitination sites. Residue contacts formed between a given lysine side chain (K2, K60 or K78) and neighboring residues, defined as those within 4 Å of any other side chain on barstar. Unfilled bars correspond to the number of residue contacts observed in the NMR structure of barstar for each lysine side chain (PDB entry 1BTA). Filled, colored bars correspond to the mean of the average number of residue contacts formed over the course of each simulation in a given condition, after removing the first 100 ns to ensure simulation equilibration. Error bars represent the standard error of the mean (*n* = 6).

**Figure S2:**
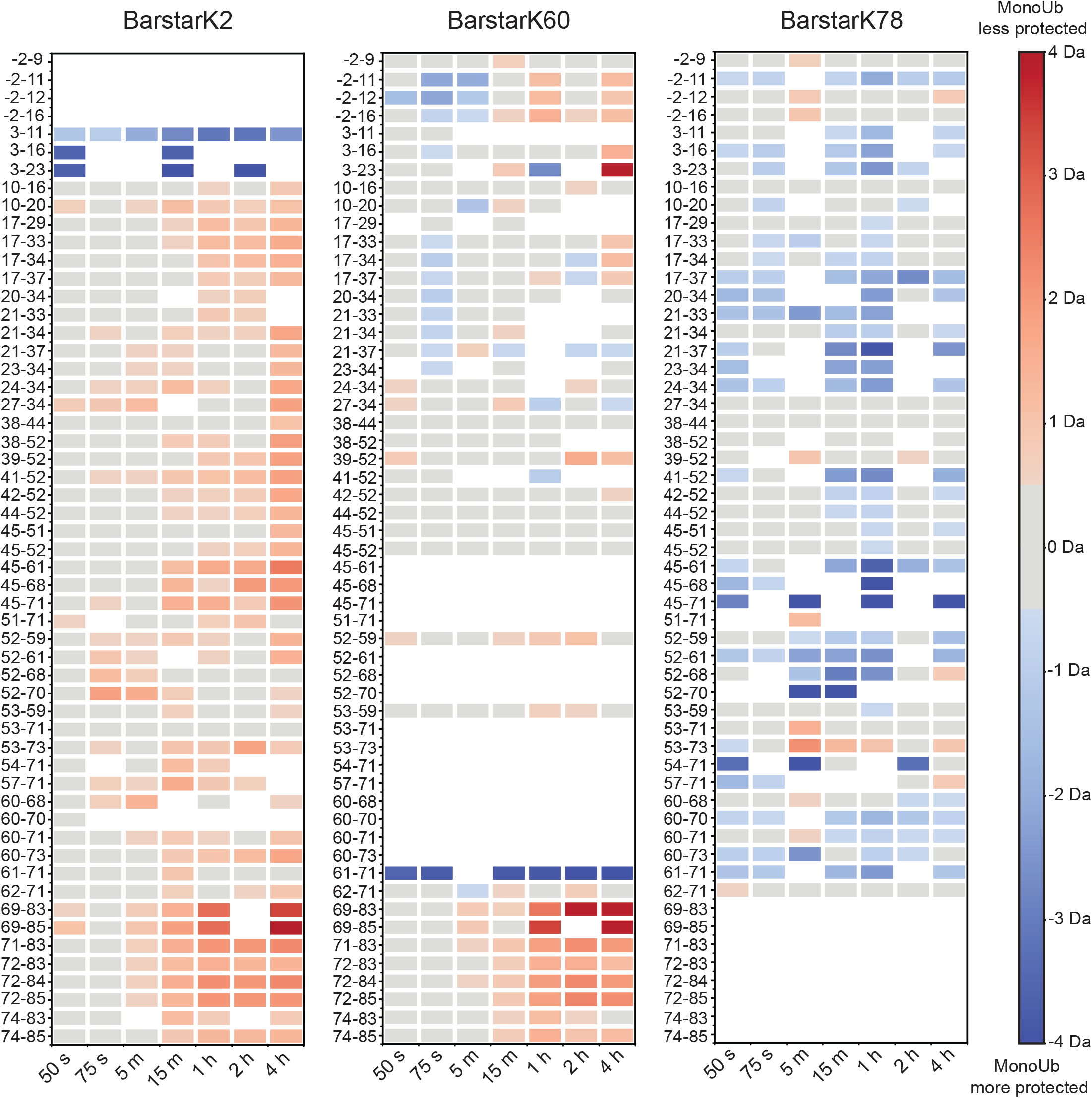
Full HDX-MS barstar peptide data set represented as difference heat maps for unmodified compared to mono-ubiquitinated barstarK2, barstarK60, and barstarK78. Heat maps showing differences in deuteration for all barstar peptides that are shared between at least two variants. Red indicates increased deuterium exchange in the mono-ubiquitinated compared to the unmodified state and blue indicates decreased deuterium uptake in the mono-ubiquitinated compared to unmodified state. We consider a deuteration difference of at least ±0.5 Da between the unmodified and mono-ubiquitinated species meaningful and use gray squares to indicate points falling below this 0.5 Da limit. In addition to the previously noted destabilization upon ubiquitination of the barstar C terminus at sensitive sites, the general destabilization observed in monoUb-barstarK2 and monoUb-barstarK60 at longer time points is consistent with the previously observed global destabilization of these mono-ubiquitinated variants. Peptides containing the site of the modification could not be analyzed because the ubiquitin ‘scar’ prevented accurate mass identification. Full peptide data sets are available as **source data**.

**Figure S3:**
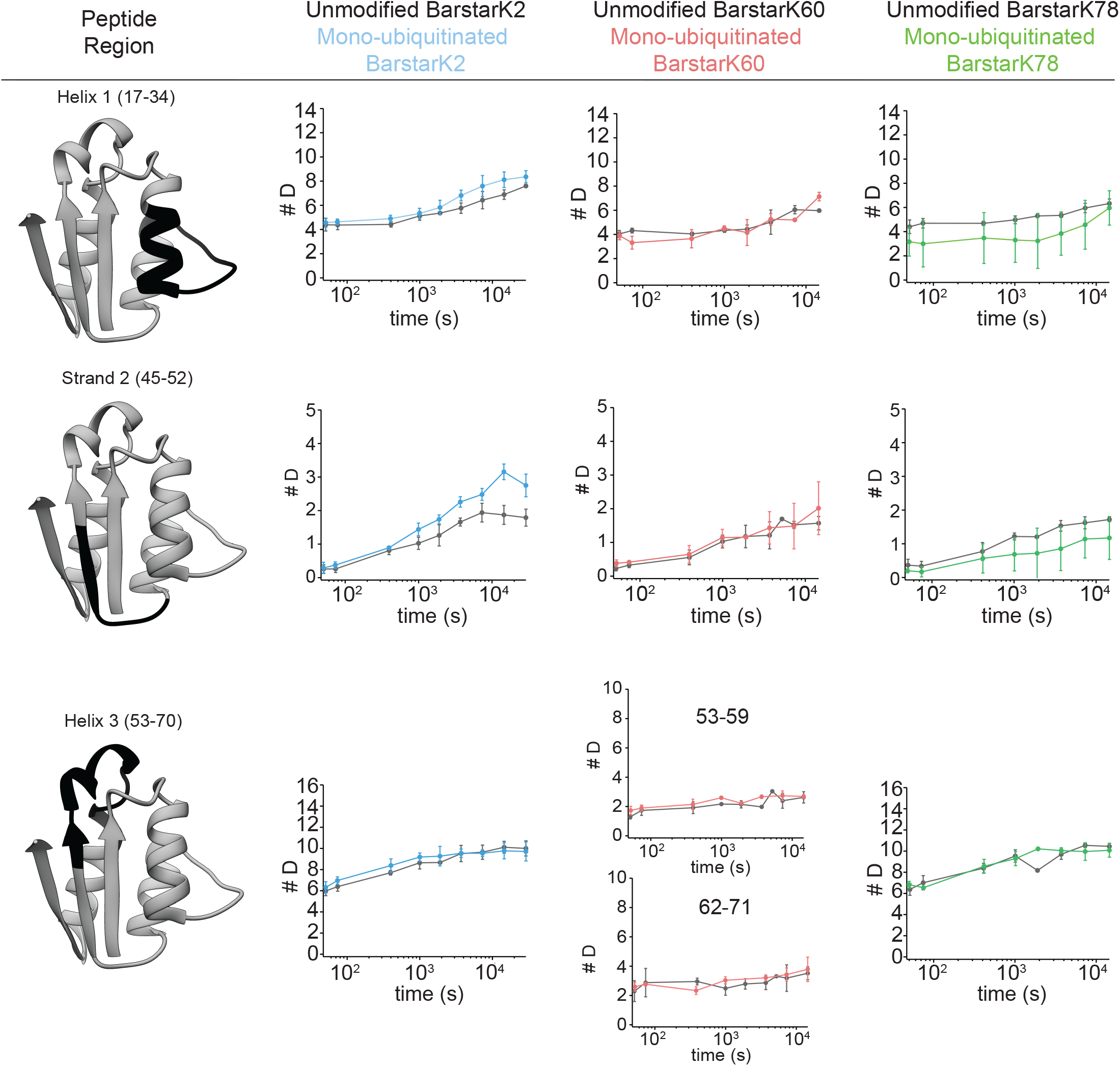
HDX-MS uptake plots for representative, non-terminal peptides. Deuterium uptake over time not corrected for back exchange for representative peptides from barstar HDX-MS experiments. Plots show unmodified (grey) and mono-ubiquitinated (blue) barstarK2 (n=3 for each protein state), unmodified (grey) and mono-ubiquitinated (red) barstarK60 (n=3 for each protein state), and unmodified (grey) and mono-ubiquitinated (green) barstarK78 (n=3 for each protein state) (PDB 1BTA). Error bars represent the standard deviation of replicates, and points are plotted at the average time that the LEAP robot performed the exchange for a given desired time point. Actual time points for each replicate are available as **source data**. Ribbon diagrams depict barstar secondary structure elements from which each peptide is derived.

**Figure S4:**
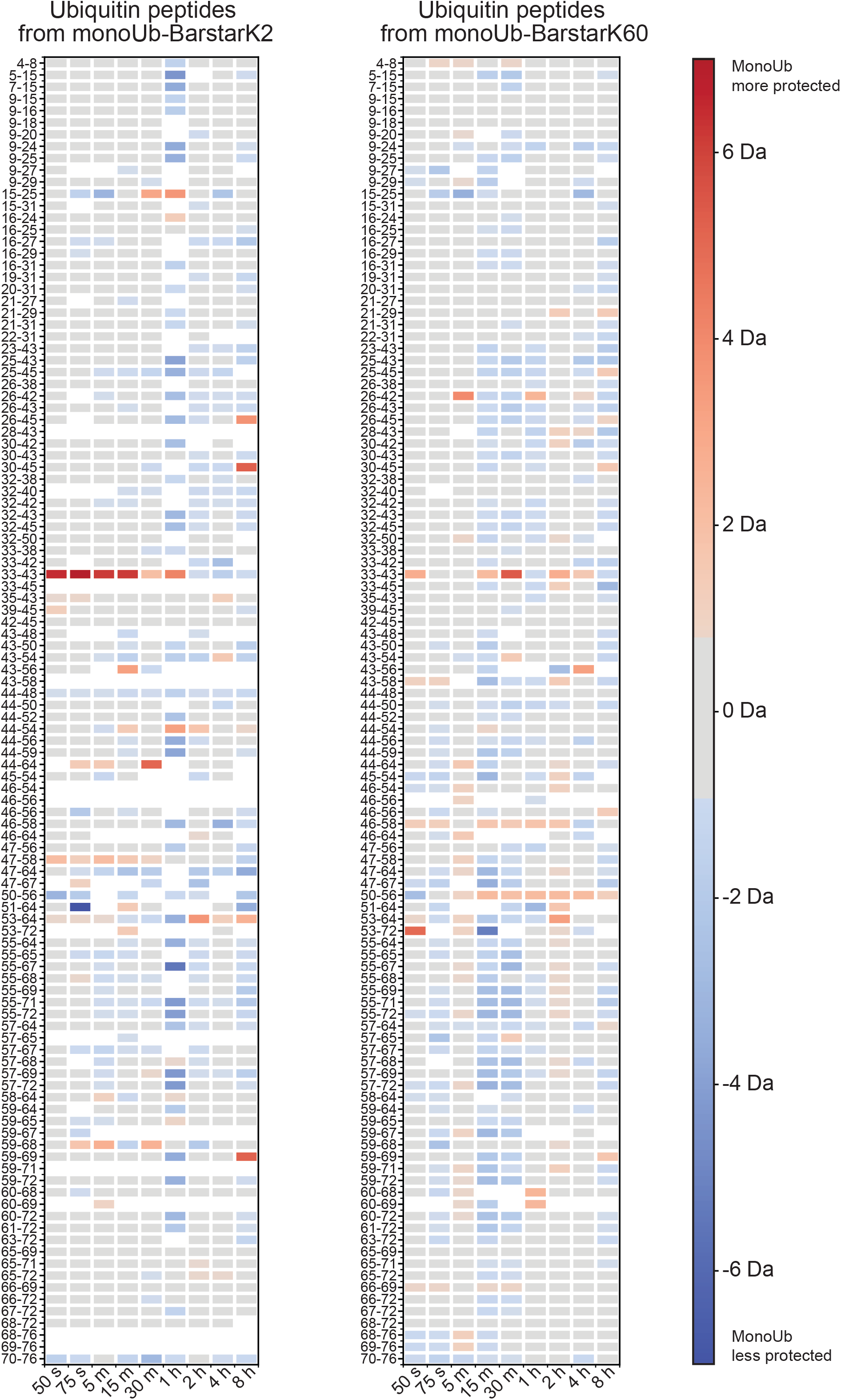
Difference heat maps ubiquitin peptides in HDX-MS experiments reveal few changes in ubiquitin upon conjugation to barstar sensitive sites. Heat maps showing differences in deuteration for unmodified ubiquitin peptides compared to peptides derived from ubiquitin when conjugated to barstarK2 and barstarK60 (the sensitive sites). Red indicates increased deuterium exchange in the barstar-conjugated compared to the unmodified state and blue indicates decreased deuterium uptake in the barstar-conjugated compared to unmodified state. We use gray squares to indicate a deuteration difference between the unmodified and mono-ubiquitinated species of less than ±0.5 Da. No notable differences are observed between unmodified ubiquitin and barstar-conjugated ubiquitin within our experimental detection limit of ~5 kcal/mol from the native state.

**Figure S5:**
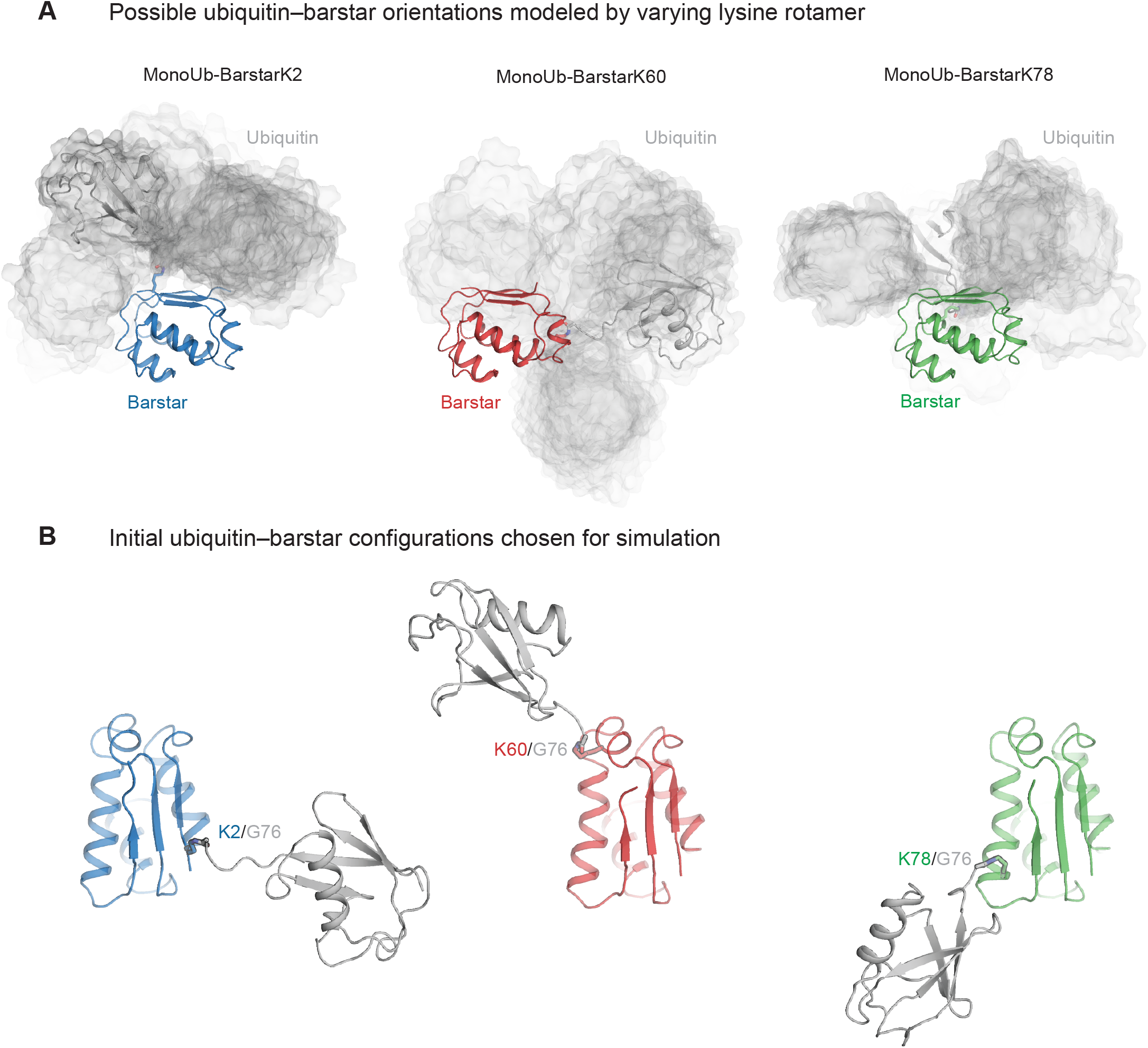
Structural models of monoubiquitinated barstar variants. **(A)** Models of barstar linked via isopeptide bond to ubiquitin at one of three sites (K2, left; K60, middle; K78, right). To generate such models, we varied the lysine rotamer at each site of attachment and show only those models lacking significant clashes between the barstar and ubiquitin surface (see Methods). Grey clouds correspond to surface representations of ubiquitin from each model (barstar not shown), overlaid after aligning each one on a structure of barstar. We show 16 models for monoUb-barstarK2, 17 models for monoUb-barstarK60, and 13 models for monoUb-barstarK78. **(B)** From each of the three sets of models shown in (A), we selected one configuration to initiate MD simulations of each variant. We selected the configurations with the maximal distance between the center of mass of barstar and the center of mass of ubiquitin.

**Figure S6:**
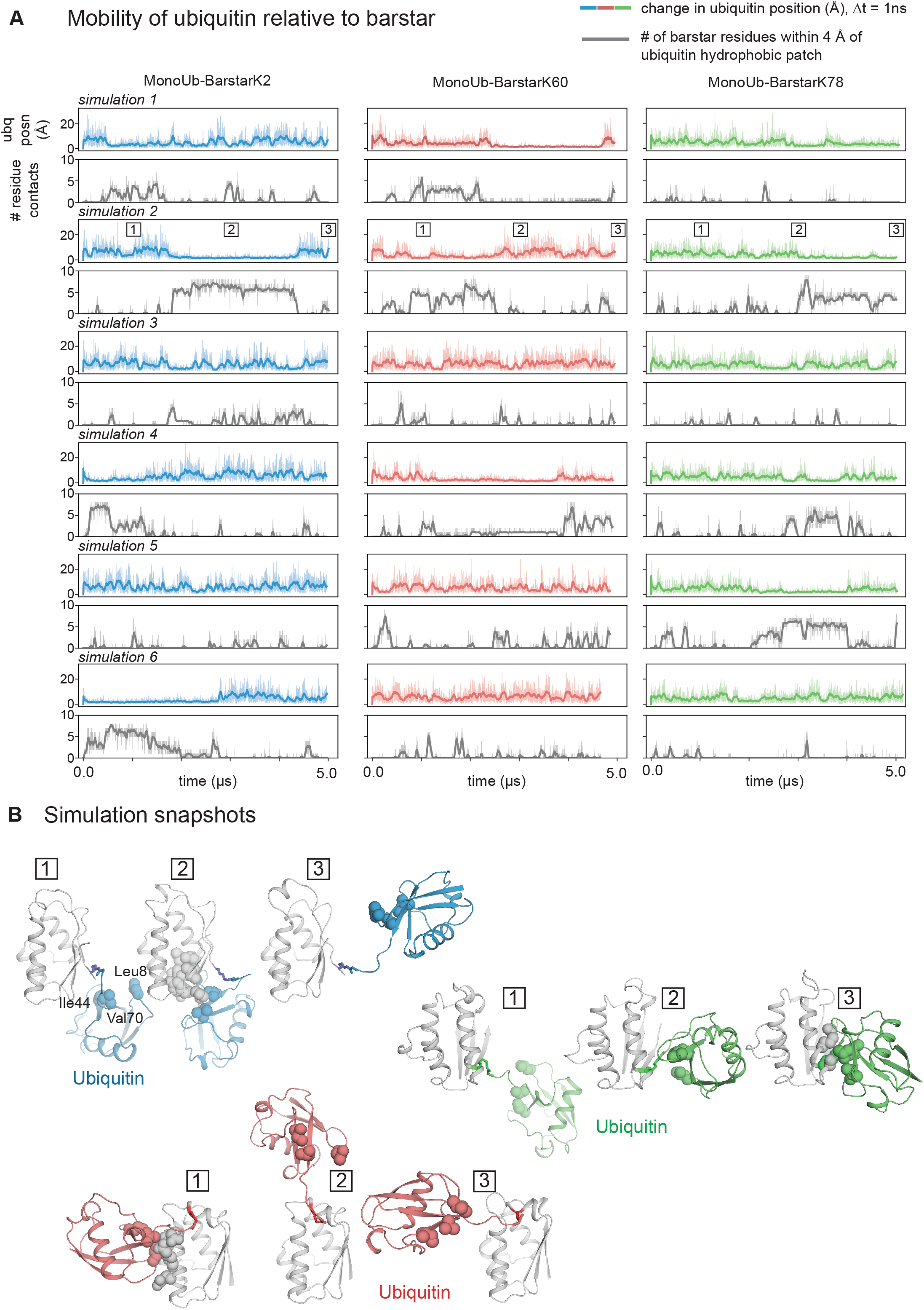
Ubiquitin mobility in simulation. **(A)** In simulation, ubiquitin typically rotates freely about the isopeptide linkage for all three monoubiquitinated barstar variants, occasionally visiting orientations that allow it to form transient surface–surface interactions with barstar. Two plots shown for each of six simulations per simulation condition. Upper plots (colors) measure the change in position of ubiquitin’s center of mass from its previous position, using a timestep of 1 ns. Lower plots (gray) show the number of barstar residues within 4 Å of a hydrophobic patch on ubiquitin (Leu8, Ile44, Val70) (85). For example, in simulation 6 of the monoUb-barstarK2 condition, ubiquitin remains close to its initial position for nearly 3 μs (blue trace; unchanging portion of plot) before becoming more mobile and visiting a range of orientations with respect to barstar (blue trace; fluctuating portion of plot). During those same 3 μs of simulation, ubiquitin and barstar form several surface–surface interactions involving this hydrophobic patch. Indeed, a reduction in ubiquitin mobility correlates with an increase in surface–surface interactions between ubiquitin and barstar (monoUb-barstarK2: *r* = −0.42; monoUb-barstarK60: *r* = −0.28; monoUb-barstarK78: *r* = −0.33, where *r* represents the Pearson correlation coefficient). Other surfaces on ubiquitin not involving the hydrophobic patch may interact with barstar’s surface, but this occurs rarely in our simulations (but see simulation 1 of the monoUb-barstarK60 condition, between 3 and 5 μs, for an example); see also **Fig. S9**. Thick traces represent a moving average, using a window of 20 ns. Thin traces represent the raw data sampled every 1 ns from each trajectory. **(B)** Simulation snapshots taken from simulation 2 as a representative of each condition reveal the high degree of mobility of ubiquitin in simulation, as well as its ability to form surface–surface interactions with barstar. Colored spheres represent the side chains of Leu8, Ile44 and Val70 of ubiquitin; gray spheres represent barstar residues within 4 Å of any of those three ubiquitin residues. Snapshots 1, 2, and 3 taken at 1.0, 3.0, and 5.0 μs, respectively.

**Figure S7:**
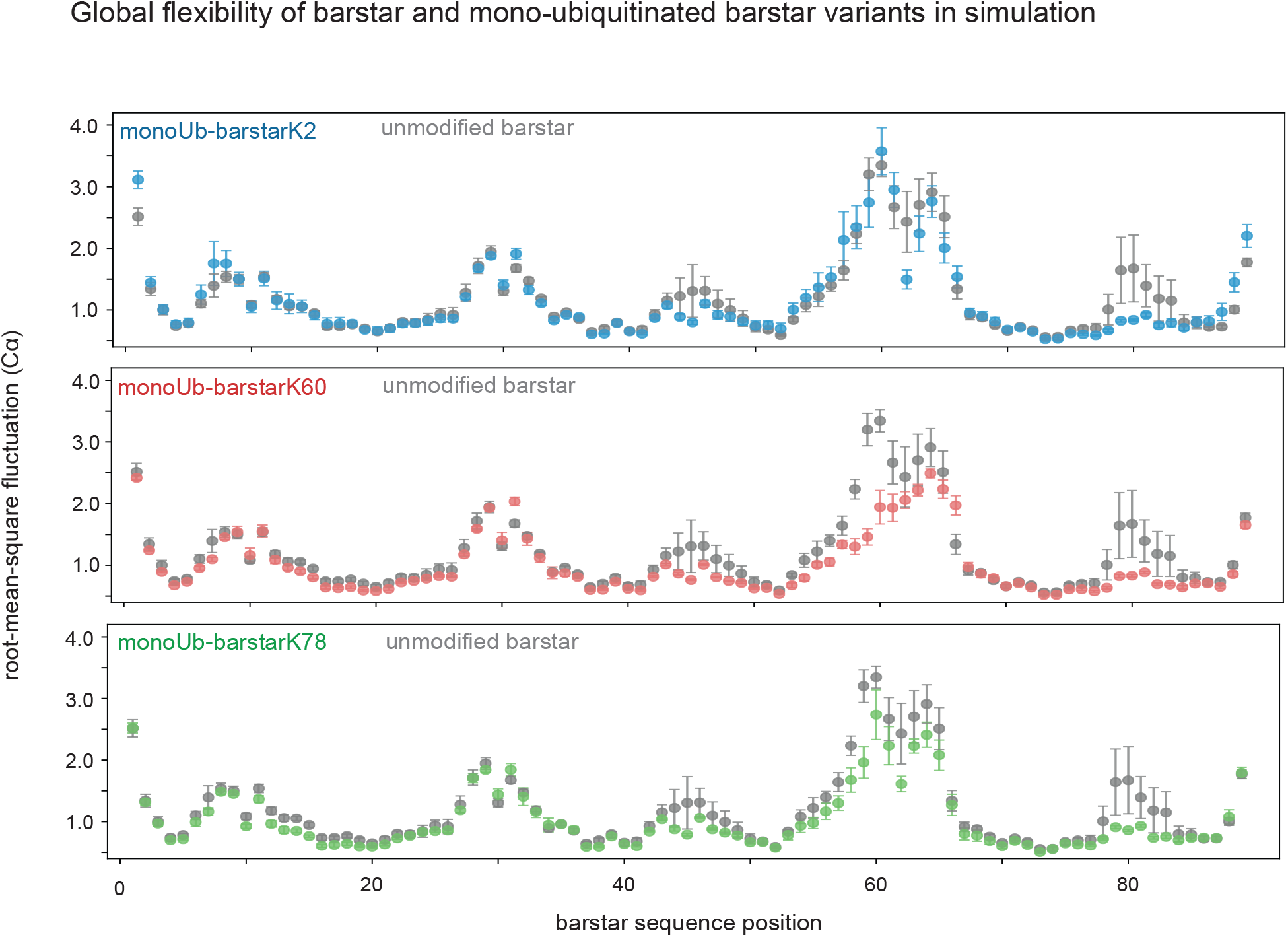
Conformational fluctuations in monoubiquitinated and unmodified barstar variants. Plots show the root-mean-square fluctuation of each Cα atom in barstar about its average position calculated for a given simulation condition, after removing the first 100 ns of each simulation to ensure equilibration. Error bars represent the standard error of the mean (*n* = 6). Plot for unmodified barstar is identical to the plot shown in **Fig. 2** and is shown here again for comparison to the monoUb-barstar conditions.

**Figure S8:**
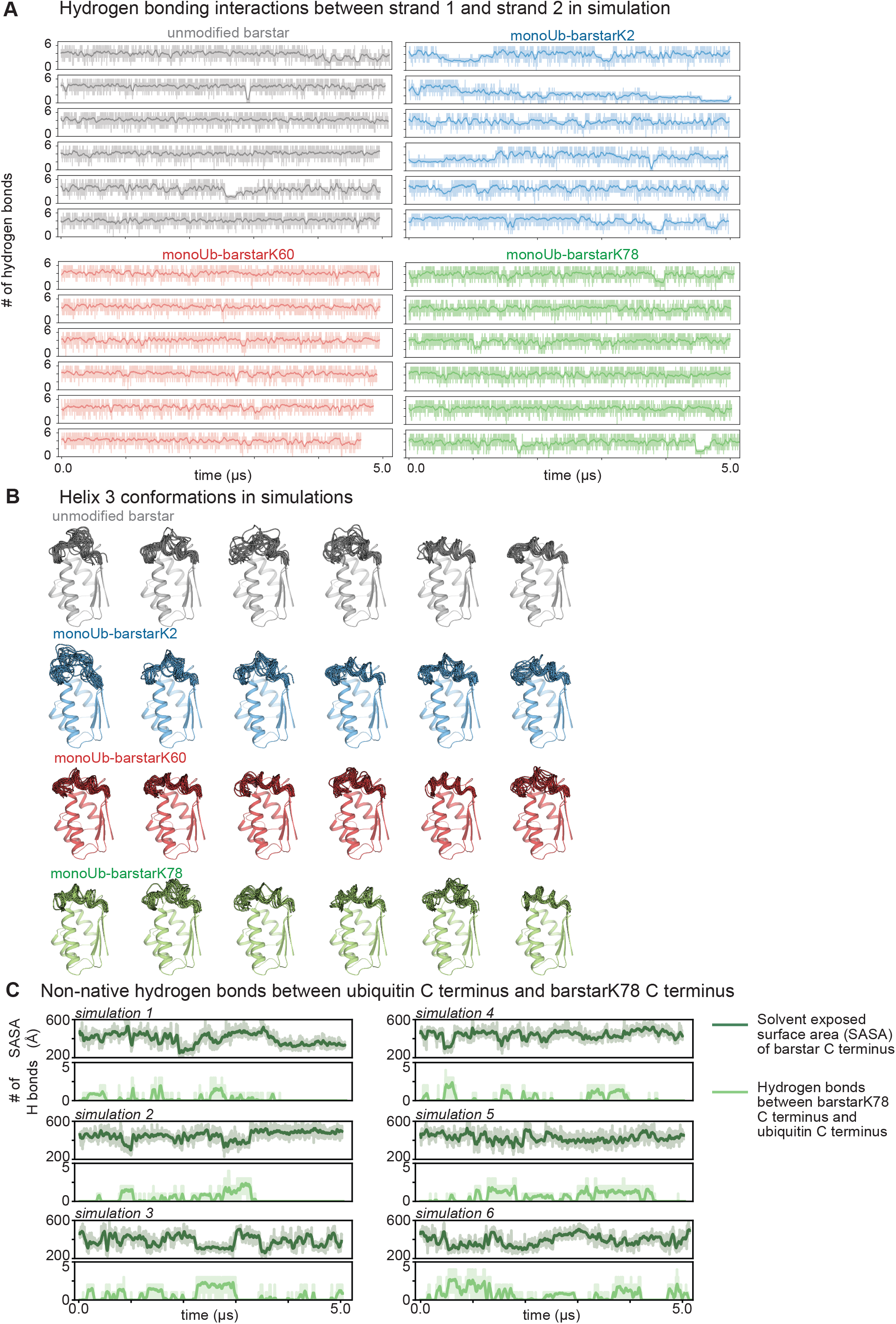
Conformational fluctuations near local sites. **(A)** Counts of hydrogen bonds between residues on the barstar N terminus (strand 1) and the neighboring strand of barstar (strand 2). Each trace corresponds to a single simulation; simulations from within one condition share the same color. In general, monoUb-barstarK2 exhibits a reduction in hydrogen bonding with strand 2 compared to other conditions. See also **Fig. 4B**. **(B)** Snapshots of unmodified barstar and the different monoubiquitinated barstar variants, sampled every 250 ns from simulation. Each structural rendering corresponds to a single simulation per condition. In general, monoUb-barstarK60 exhibits the least amount of fluctuation, and the greatest of helicity, in helix 3, the helix containing K60. See also **Fig. 4A**. **(C)** Counts of hydrogen bonds formed between residues on the ubiquitin C terminus and the barstarK78 C terminus (one plot shown per simulation). Hydrogen bond formation occurs spontaneously and frequently across multiple simulations. Formation of multiple hydrogen bonds correlates with a decrease in the accessible surface area of the barstar C terminus (residues 83–89; see Methods), suggesting that the barstarK78 C terminus becomes protected upon hydrogen bonding to the ubiquitin C terminus.

**Figure S9:**
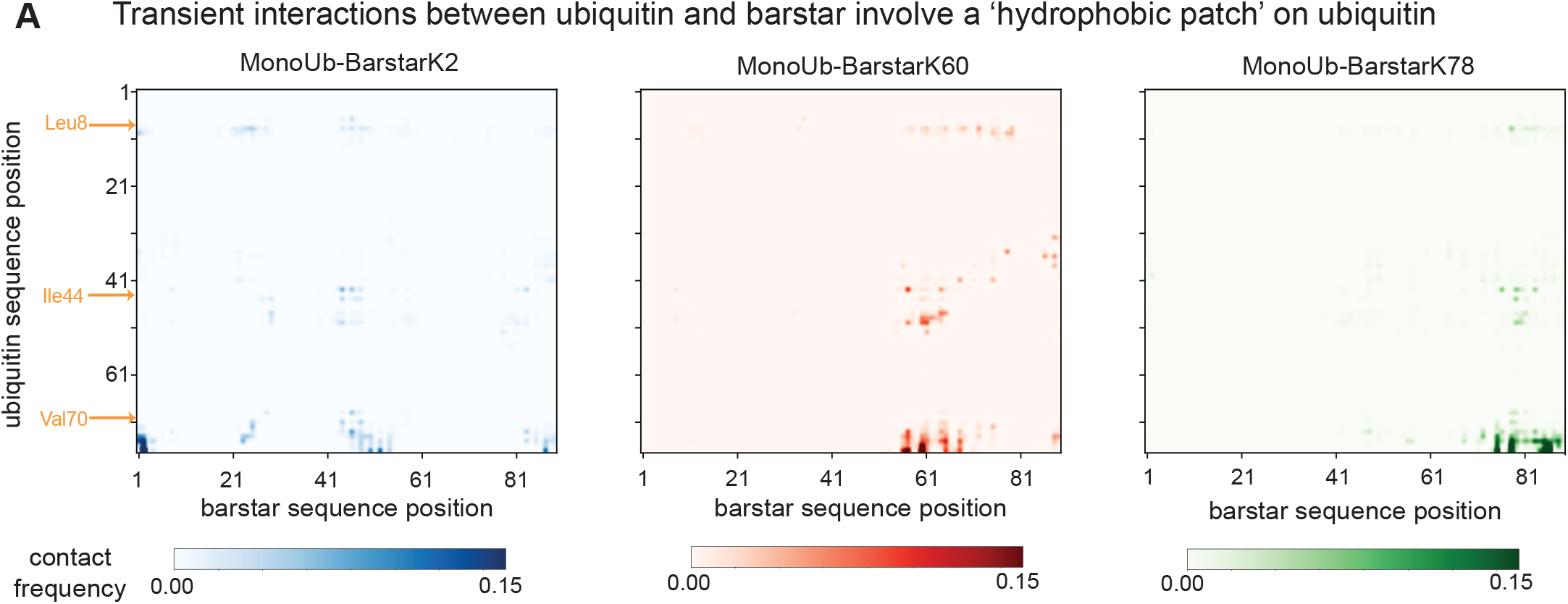
Contact maps for ubiquitin–barstar interactions. We calculated the frequency of contacts formed between residues on ubiquitin and residues on barstar across all six simulations per simulation condition (see Methods). These plots demonstrate that (1) frequency of interaction between residues on ubiquitin and those on barstar is quite low for any residue– residue pair, as indicated by the color bars, and (2) when surface–surface interactions do occur, they tend to involve residues in the vicinity of the Ile44 hydrophobic patch, described in **Fig. S6**. Hydrophobic patch residues Leu8, Ile44 and Val70 are highlighted in orange.

**Figure S10:**
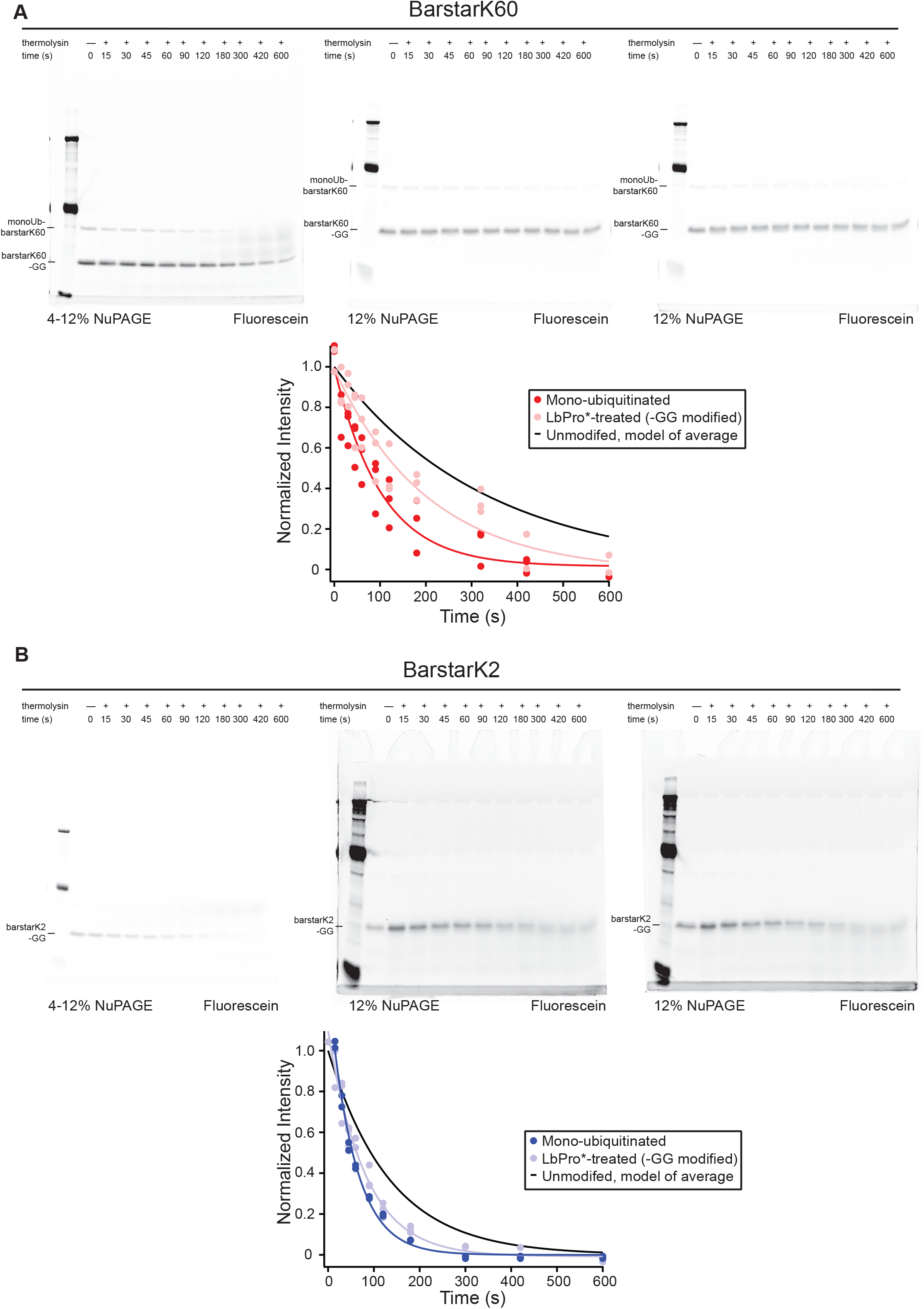
Native-state proteolysis experiments on LbPro*-treated mono-ubiquitinated barstarK2 and barstarK60 reveal that ubiquitin ‘bulk’ is important for modulating destabilization. **(A)** Gels and gel quantifications for native-state proteolysis experiments on LbPro*-treated mono-ubiquitinated barstarK60. BarstarK60 is labeled on its single cysteine, C82, with fluorescein-maleimide and gels are imaged in Typhoon fluorescein channel. Proteolysis experiments were performed using 0.2mg/mL thermolysin protease. Band intensities are normalized to the no protease condition and then all replicates are fit simultaneously to a first order exponential (dark red = mono-ubiquitinated band n = 3, light red = LbPro*-treated/barstarK60 –GG band n = 3). The exponential model previously calculated for unmodified barstarK60 is plotted in black. The intermediate proteolysis rate of LbPro*-treated barstarK60 (faster than unmodified barstarK60 but slower than mono-ubiquitinated barstarK60) indicates that the full mass of ubiquitin is necessary to impart the full magnitude of the energetic destabilization. **(B)** Gels and gel quantifications for native-state proteolysis experiments on LbPro*-treated mono-ubiquitinated barstarK2. BarstarK2 is labeled on its single cysteine, C82, with fluorescein-maleimide and gels are imaged in Typhoon fluorescein channel. Proteolysis experiments were performed using 0.2 mg/mL thermolysin protease. Band intensities are normalized to the no protease condition for replicate 1 (left) and to the 15s time point for replicates 2 and 3 (middle and right) due to a gel loading error for the no protease condition. The LbPro* reaction for replicate 1 went to completion such that no mono-ubiquitinated barstarK2 band remained for quantification. All replicates are fit simultaneously to a first order exponential (dark blue = mono-ubiquitinated band n = 2, light blue = LbPro*-treated/barstarK2 – GG band n = 3). The exponential model previously calculated for unmodified barstarK2 is plotted in black. The intermediate proteolysis rate of LbPro*-treated barstarK60 (faster than unmodified barstarK2 but slower than mono-ubiquitinated barstarK60) indicates that the full mass of ubiquitin is necessary to impart the full magnitude of the energetic destabilization.

**Figure S11:**
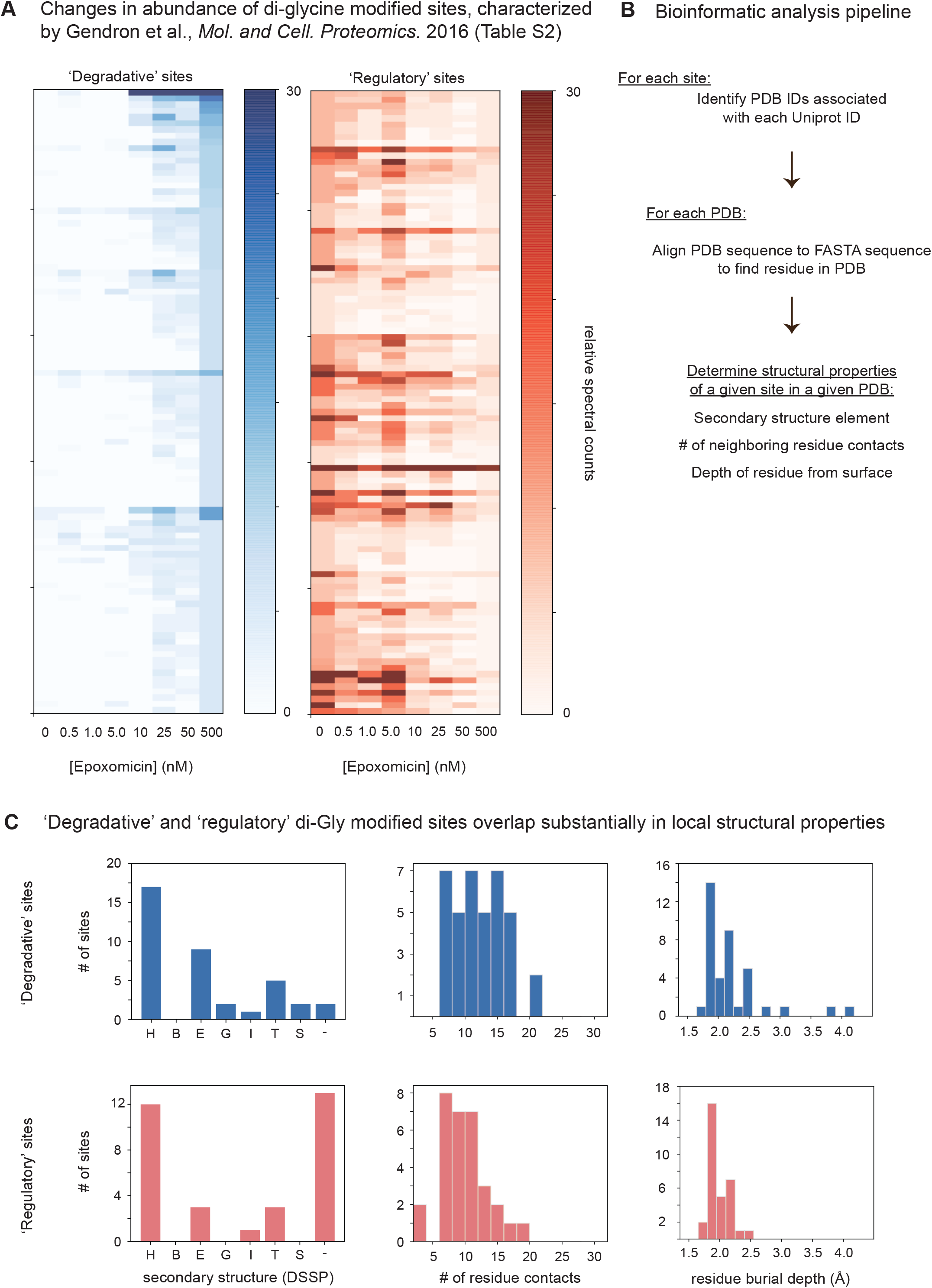
Analysis of ubiquitination sites identified through a proteomics screen in (56). **(A)** Reproduction of data displayed in Table S2 of Gendron et al. (56). Heatmaps show spectral counts of di-glycine modified sites, indicative of ubiquitination, for a proteomics screen using different concentrations of a proteasome inhibitor. Spectral counts are relative, because they are normalized by the site yielding the maximum spectral count at a particular concentration. ‘Degradative’ sites—i.e. sites whose ubiquitination leads to degradation by the proteasome—increase in abundance with proteasome inhibition, while ‘regulatory’ sites—sites whose ubiquitination has an effect on regulation or signaling—decrease in abundance with proteasome inhibition. Several caveats are associated with these classifications, however. For example, pools of ubiquitin-modified substrates can vary depending on the type of inhibitor used, and other non-ubiquitination modifications can also yield di-Gly modified lysine residues after trypsin digest (86). **(B)** Bioinformatics pipeline for extracting structural information about di-glycine modified ubiquitination sites. See Methods for details. **(C)** Results of structural analyses performed on PDB structures containing sites of interest shown in (A). Note that PDB structures containing such sites exist for 70 (i.e. <50%) of these sites. Left: secondary structure analysis of the site of di-glycine modification reveals that degradative and regulatory sites occur in various regions of secondary structure. Classifications from DSSP (82): ‘H’ −4-12 alpha helix; ‘B’ - isolated beta-bridge residue; ‘E’ - strand; ‘G’ - 3-10 helix; ‘I’ - pi helix; ‘T’ - turn; ‘S’ - bend; ‘-’ - none. Reported secondary structure values for a given site correspond to the most common secondary structure elements based on all structures analyzed. The reported number of residue contacts, as well as residue burial depth, were determined based on a single representative PDB structure for each site. See Methods for details.

